# Redox-regulated cysteine acylation governs β-lactam sensing by the *Vibrio* histidine kinase VbrK

**DOI:** 10.64898/2026.06.08.730885

**Authors:** Ignacio Guido Palanca, Irina Paula Suárez, Maria Jose Marcaida, Luciano Andres Abriata, Natalia Gasilova, Laure Menin, Franco Emanuel Lacava, Julieta Cairoli, Matteo Dal Peraro, Leticia Irene Llarrull

**Author notes:** Corresponding author/s: Name: Leticia Irene Llarrull, Mailing Address: Instituto de Biología Molecular y Celular de Rosario (IBR-CONICET-UNR), Ocampo y Esmeralda, Edificio IBR, Predio CONICET, Rosario, 2000, Santa Fe, Argentina, Interno 638, Name: Irina Paula Suárez, Mailing Address: Instituto de Biología Molecular y Celular de Rosario (IBR-CONICET-UNR), Ocampo y Esmeralda, Edificio IBR, Predio CONICET, Rosario, 2000, Santa Fe, Argentina Interno 638. These authors contributed equally to this work (Ignacio Guido Palanca, Irina Paula Suárez). Keclon S.A., Cerrito 847, 2000 Rosario, Argentina.

## Abstract

The ability of pathogens to develop resistance mechanisms makes the continuous search for novel therapeutic targets indispensable. Resistance-activating systems are promising targets for restoring the efficacy of existing antibiotics. VbrK/VbrR is a two-component system from *Vibrio parahaemolyticus* reported as the first sensing system in gram-negatives that directly detects β-lactam antibiotics. β-Lactam-induced activation of this system results in the expression of the serine β-lactamase CARB. In this study, we provide insights into the mechanism of β-lactam binding to the periplasmic sensor domain of the histidine kinase VbrK. Our results demonstrate that the interaction depends on the redox state of cysteines C86 and C107, highlighting the role of disulfide bond dynamics in modulating ligand recognition. We further show that formation of a non-covalent complex leads to acylation of the sensor domain by β-lactams, a modification that is slowly reversed through de-acylation, yielding the hydrolyzed β-lactam ring and allowing for recovery from induction once the antibiotic has been depleted from the environment. Together, these findings reveal a previously unrecognized redox- and covalent chemistry–dependent mode of β-lactam interaction with histidine kinases, providing a molecular framework to understand how VbrK detects and responds to β-lactam antibiotics, and opening new avenues to prevent manifestation of resistance.

**Short broader audience statement:** Bacterial pathogens can sense the presence of antibiotics and trigger resistance responses, making infections increasingly difficult to treat. In this study, we uncover a previously unknown mechanism by which β-lactam antibiotics interact with a sensor protein that activates resistance in *Vibrio parahaemolyticus*. These findings pave the way for designing new compounds that can block this interaction and thus restore the effectiveness of β-lactam antibiotics against gastrointestinal infections caused by resistant bacteria.

## Introduction

*Vibrio parahaemolyticus* is one of the most common causes of gastroenteritis from consuming contaminated seafood worldwide^1–4^. Infections from this pathogen can be fatal in immunocompromised patients or those with debilitating medical conditions like liver failure or diabetes^5^. Most clinical and environmental isolates of *V. parahaemolyticus* show resistance to β-lactam antibiotics, which limits treatment options. Essentially all *V. parahaemolyticus* isolates encode a chromosomal class A serine β-lactamase (CARB; *bla*_V110_ or *vpa0477* gene) that efficiently hydrolyzes carbenicillin and other penicillins^6^. The expression of this serine β-lactamase is induced in the presence of β-lactams by the VbrK/VbrR system^7^.

VbrK (**V**ibrio **β**-lactam **r**esistance sensor **k**inase) is a transmembrane sensor histidine kinase that contains a globular periplasmic domain (VbrK^SD^ hereafter for “VbrK Sensor **D**omain”) at its N-terminus. VbrK undergoes autophosphorylation in *V. parahaemolyticus* in the presence of β-lactams^7^. It has been proposed that the interaction of the β-lactams with the periplasmic domain of VbrK triggers the signaling cascade, via the response regulator VbrR, that leads to transcription of the *bla*_*V110*_ gene and the regulation of a master regulon ExsACD which leads to the repression of a type 3 secretion system which regulates the virulence of the pathogen^8^. Interestingly, the VbrK/VbrR system has also been shown to respond to S-nitrosylation of residue cysteine 86, one of the four cysteine residues located in VbrK^SD 8^. The C86S substitution in VbrK eliminates its nitrite-induced autophosphorylation and abolishes nitrite-mediated repression of the type 3 secretion system, thereby attenuating pathogen virulence.

Several crystallographic structures of VbrK^SD^ have been solved, including homologs from *Vibrio cholerae* and *V. rotiferianus*^*9–12*^, where two subdomains can be observed, one proximal and the other distal to the membrane, while the N- and C-termini lie in close proximity. Each subdomain contains a disulfide bridge: C86–C107 and C226–C234 in the distal and proximal subdomains, respectively. The mechanism of activation of VbrK has not been elucidated so far. In fact, there is controversy in the literature on the nature of *β*-lactam binding to the kinase. Direct interaction of penicillin G with VbrK was demonstrated by Zhou and collaborators, using an experimental set up to detect penicillin binding to purified and immobilized VbrK^SD^ with an anti-penicillin antibody^7^. In contrast, no interaction was evidenced between VbrK^SD^ and Penicillin G in ITC experiments by Lescar and collaborators^10^. To date, no antibiotic-bound VbrK^SD^ structure has been reported.

Here, we confirmed that β-lactam antibiotics bind to VbrK^SD^ and we documented the modulation of this interaction by the redox state of cysteines C86 and C107. We reveal the formation of a covalent adduct of β-lactam antibiotics with the isolated sensor domain of VbrK and with full-length VbrK. Combining NMR experiments, labeling experiments, mass spectrometry and stopped-flow kinetic assays, we unveiled the reaction mechanism: formation of a non-covalent β-lactam:VbrK^SD^ adduct, followed by formation of a stable covalent complex, and very slow release of the hydrolyzed antibiotic. Covalent modification of the sensor domain of VbrK is due to formation of β-lactam–derived thioesters, which we propose stabilize the activated state of the sensor histidine kinase, thereby driving β-lactam–induced activation of the VbrK/VbrR signaling pathway. These findings reveal a previously unrecognized redox- and covalent chemistry–dependent mode of β-lactam interaction with histidine kinases, providing a molecular framework to understand how VbrK detects and responds to β-lactam antibiotics.

## Results

### VbrK^SD^ interacts with β-lactams when cysteines are reduced and is selective for the intact β-lactam ring

To explore how the VbrK sensor domain interacts with β-lactams, we optimized its expression and purification, confirming the presence of an N-terminal signal peptide (residues 1-22) that was cleaved to yield the mature version of the protein (Figures S1). The same cleavage site was identified in the isolated sensor domain and in the full-length protein (Figure S2).

The X-Ray structures of VbrK^SD^ from *Vibrio parahaemolyticus* (PDB IDs: 7CUS and 7CJR) show the presence of two disulfide bonds (C86–C107 and C226–C234). It has been documented that VbrK phosphorylation in response to nitrite is due to S-nitrosylation of cysteine residue 86^8–10^, a reaction that would require the reduction of the C86–C107 disulfide bond. We determined the redox state of cysteines in purified VbrK^SD^ using Ellman’s reaction^13^. Freshly purified VbrK^SD^ contained no reduced cysteine residue, whereas TCEP-treated VbrK^SD^ exhibited two reduced cysteines (Figure S3). Secondary structure conservation after this treatment was confirmed by circular dichroism (Figure S4), and conservation of the three-dimensional structure was confirmed in ^1^H-^15^N-HSQC NMR experiments (Figure S5).

Saturation transfer difference ^1^H Nuclear Magnetic Resonance (STD NMR) experiments^14^ were employed to assess the interaction of VbrK^SD^ with antibiotics and whether the interaction was dependent on the redox state of cysteine residues. STD NMR spectroscopy is especially suited for the study of low to medium affinity complexes, resulting in a difference spectrum where the signals observed indicate that the saturated atoms of the protein were close enough in space to those of the ligand for Nuclear Overhauser Effect (NOE) to occur. STD NMR experiments were conducted on ampicillin solutions in the presence of freshly purified VbrK^SD^ (no reduced cysteine residue, from now onwards referred to as oxidized) or TCEP-treated VbrK^SD^ (with one reduced disulfide bond) (**Figure 1**). ^1^H chemical shifts assignment for ampicillin in aqueous solution were taken from Upadhyay *et al*.^*15*^ The STD NMR experiment with oxidized VbrK^SD^ (no reduced cysteine) showed no signals corresponding to ampicillin protons (**Figure 1** and Figure S6). However, signals of ampicillin protons at 7.37, 5.39, 4.10 ppm (**Figure 1**.A) and 1.42 and 1.35 ppm (**Figure 1**.B) were detected in experiments with TCEP-incubated protein, evidencing the perturbation of these protons due to the spatial proximity of the saturated protons on the protein. The STD NMR experiments demonstrated unequivocally the non-covalent interaction of ampicillin with the VbrK^SD^ sample exhibiting two reduced cysteine residues. Based on the characteristics of the STD NMR experiment, ampicillin binding to reduced VbrK^SD^ is reversible, with moderate to low affinity. All ampicillin signals detected in the ^1^H NMR spectrum were perturbed by saturation transfer (squares in **Figure 1**.A and B, color coded in **Figure 1**.C), indicating that the entire molecule engaged in the interaction with VbrK^SD^. We then blocked the reduced cysteines with iodo-acetamide (IAA) and repeated the STD NMR experiment with ampicillin (**Figure 1**.A and B). None of the previously detected signals of ligand protons were observed, indicating that the IAA-modified protein did not interact with ampicillin, thereby suggesting that the reduced cysteines are essential for β-lactam binding. Next, we tested the selectivity of the interaction by acquiring an STD NMR spectrum on a sample of reduced VbrK^SD^ in the presence of a mixture of 400 µM fresh ampicillin and 400 µM hydrolyzed ampicillin. The experiment showed that only the protons on the methyl groups of fresh (non-hydrolyzed) ampicillin were perturbed by the interaction with the protein (**Figure 1**.D), indicating that the binding site discriminates between the hydrolyzed and non-hydrolyzed β-lactam ring.

**Figure 1.**
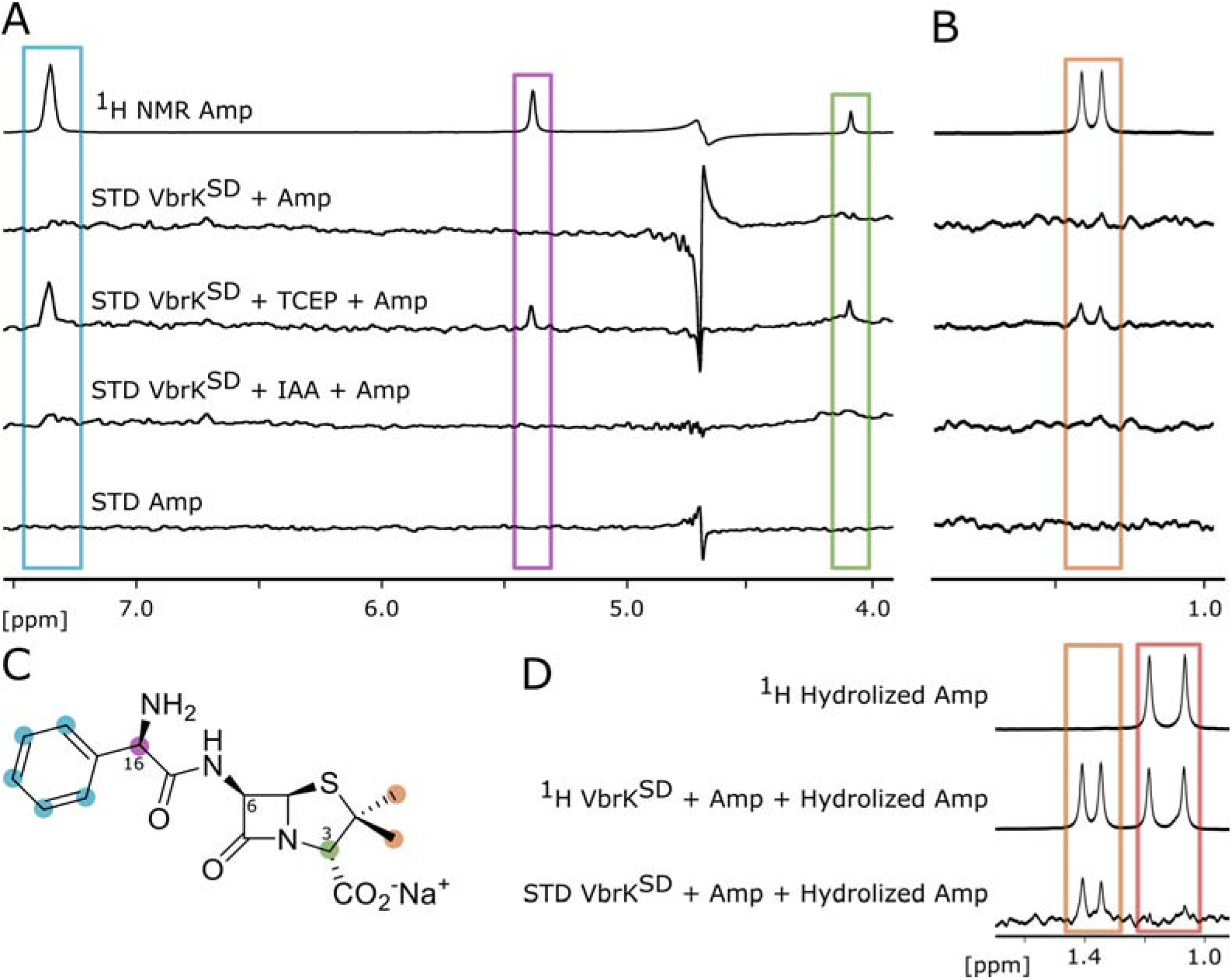
VbrK^SD^ interacts with ampicillin when two cysteines are reduced. **A**. STD NMR experiments acquired on 400 µM ampicillin (Amp) with 20 µM oxidized, reduced or IAA-treated VbrK^SD^ and on 400 µM Amp without protein. The first spectrum (Top) corresponds to the ^1^H NMR spectrum of 400 µM Amp; successive lower spectra correspond to STD NMR spectra saturating at 0.75 ppm. **B**. Same samples as in A, but STD NMR experiment saturating at 7 ppm. In panels A and B, the colored rectangles indicate the ampicillin proton corresponding to each signal, as highlighted in **C. C**. Ampicillin structure showing the color-code for each proton resonance signal. **D**. Top spectrum corresponds to ^1^H NMR spectrum of 400 µM NDM1-hydrolyzed Amp. Middle spectrum corresponds to ^1^H NMR of 400 µM Amp + 400 µM NDM1-hydrolyzed Amp + 20 µM TCEP-treated VbrK^SD^. Bottom spectrum corresponds to STD NMR spectrum acquired on the same sample, saturating at 7 ppm.

After establishing that detection of β-lactam binding required treatment of VbrK^SD^ with a reducing agent, we investigated which β-lactam families were recognized by VbrK and could therefore potentially activate β-lactamase expression in *Vibrio*. We evaluated the interaction of TCEP-treated VbrK^SD^ with the penicillins oxacillin and penicillin G, the cephalosporin cefotaxime and the carbapenem meropenem (**Figure 2**). The detection of antibiotic proton signals in the STD NMR experiments with penicillin G and oxacillin indicated binding to VbrK^SD^. As with ampicillin, the STD NMR spectra presented signals corresponding to all penicillin G and oxacillin proton resonances, suggesting that the entire antibiotic molecules were engaged in interaction within VbrK^SD^. In the case of cefotaxime, fewer signals were perturbed, suggesting that its structural differences from penicillins translate into a distinct mode of interaction with VbrK^SD^. Finally, no STD signal was detected with meropenem, suggesting carbapenems might not be detected by this system. This interaction profile correlates with the substrate spectrum of the CARB β-lactamase from *V. parahaemolyticus*, which is active against penicillins and some cephalosporins (nitrocefin, cefepime and cefpirome) but does not hydrolyze cefotaxime, cefuroxime nor the carbapenem imipenem^6,7^.

**Figure 2.**
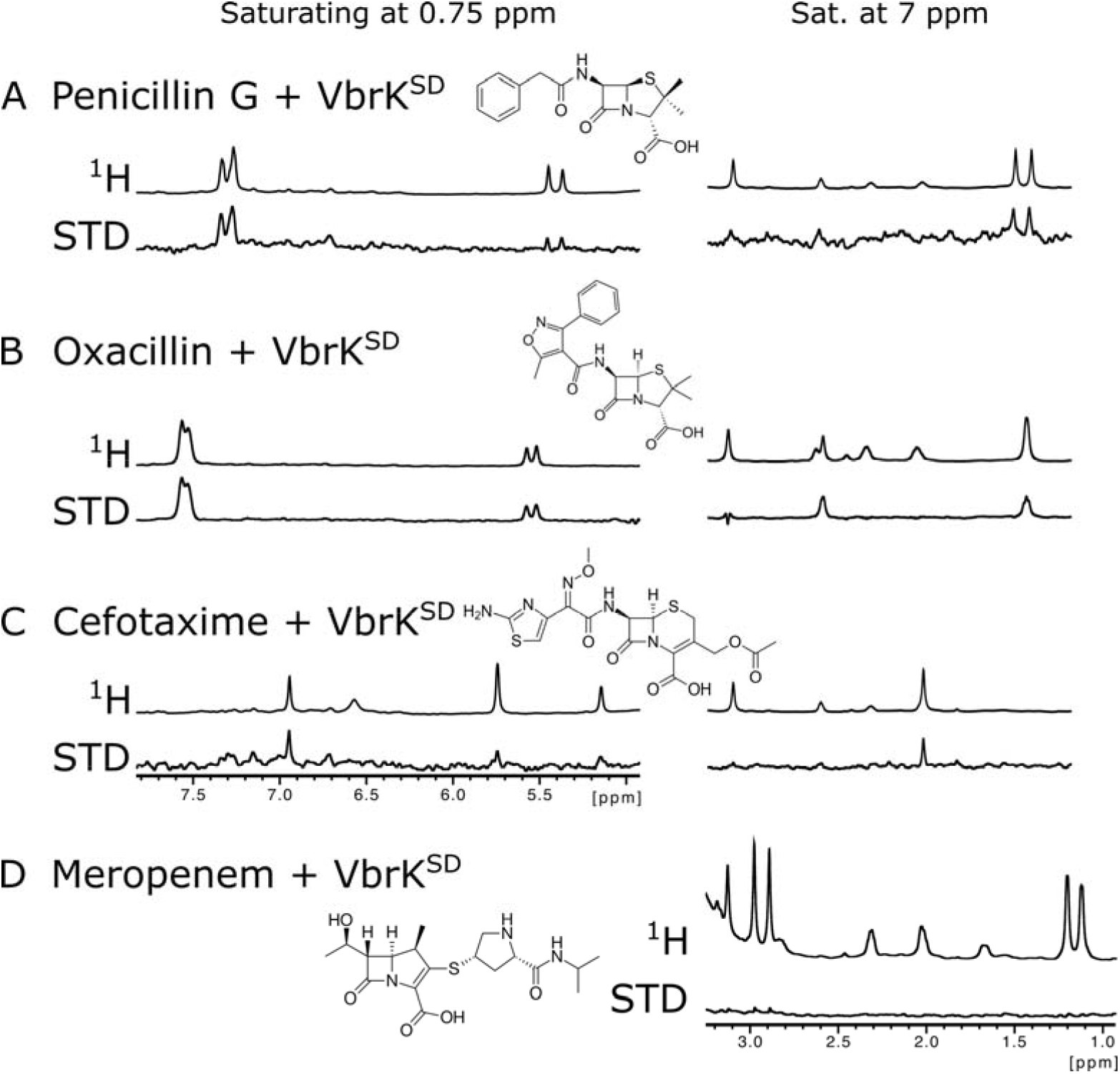
VbrK^SD^ interacts with penicillins and cephalosporins, but not with carbapenems. A 20 µM sample of TCEP-treated VbrK^SD^ was mixed with 400 µM of: **A**. penicillin G; **B**. oxacillin; **C**. cefotaxime; **D**. meropenem. In each panel, the top spectrum corresponds to the ^1^H spectrum of the antibiotic and the bottom spectrum corresponds to the STD NMR assay. The STD NMR spectra displayed on the left were acquired saturating at 0.75 ppm and those on the right, saturating at 7 ppm.

As controls for these experiments, we tested interaction of the reduced domain with two different ligands that undergo unrelated biology, the glycopeptide vancomycin (a non-β-lactam antibiotic) and the carbohydrate arabinose. None of these molecules showed difference peaks in the STD NMR spectra (Figure S7), highlighting the specificity of the protein for β-lactams.

### The VbrK^SD^ C86-C107 disulfide bond is reduced by TCEP treatment

VbrK^SD^ has four cysteines in its sequence, which are engaged in two disulfide bridges in the crystallographic structures, C86-C107 and C226-C234. The VbrK/VbrR system is activated upon S-nitrosylation of residue cysteine 86^8^. Hence, we wondered whether the disulfide bridge being reduced upon TCEP-treatment, and potentially allowing for β-lactam recognition, could be the one formed by C86 and C107. To test this, we expressed and purified the VbrK^SD^ mutants C86A, C107A, and C86A-C107A. CD spectra and the distribution of peaks in ^1^H-^15^N HSQC spectra indicated proper folding of all three mutants (Figure S4 and S5).

We quantified reduced cysteines using Ellman’s reaction on freshly purified samples and after treatment with TCEP. The analysis revealed less than one reduced cysteine per molecule in freshly purified C86A and C107A, while both TCEP-treated samples presented nearly one reduced cysteine per protein molecule (Figure S3). TCEP can reverse many common cysteine modifications, such as S-nitrosylation, S-sulfenylation, S-glutathionylation and disulfide bonds^16^. Hence, the freshly purified single mutants C86A and C107A samples contained a major fraction with a reduced cysteine per protein molecule and a small fraction of molecules where the free cysteine is blocked, most likely by modifications that could be reversed by TCEP treatment. No reduced cysteines were detected in the TCEP-treated double mutant VbrK^SD^ C86A-C107A (Figure S3). This indicated that the two cysteines being reduced upon TCEP treatment on the wild-type sensor domain were C86 and C107. This disulfide bridge is located on the VbrK^SD^ distal subdomain, the region which has been proposed to contain the β-lactam binding site^7^.

To corroborate that the two cysteines being reduced upon TCEP treatment on the wild-type sensor domain were C86 and C107, the ^1^H-^15^N HSQC spectra of untreated and TCEP-treated VbrK^SD^ were compared with those of the C86A and C107A mutants. The spectra of wild-type VbrK^SD^ showed changes in the position of some cross-peaks upon TCEP treatment (**Figure 3**.A), while the ^1^H-^15^N HSQC spectra of TCEP-treated wild-type VbrK^SD^ and the C86A and C107A mutants overlapped largely (**Figure 3**.B-C), supporting our proposal that the C86-C107 disulfide bond was the one reduced in the wild-type sensor domain of VbrK. Intact protein mass spectrometry further confirmed these observations. The mass difference between oxidized and TCEP-treated VbrK^SD^ was consistent with reduction of a single disulfide bond, corresponding to the addition of two protons upon reduction (Figure S1). Furthermore, the mass differences between TCEP-treated VbrK^SD^ and the C107A mutant, and between TCEP-treated VbrK^SD^ C107A and the C86A/C107A double mutant, were in excellent agreement with the expected mass change associated with Cys-to-Ala substitutions (Figures S10 and S12). These data indicated that the C226–C234 disulfide bond remained closed under these reducing conditions.

**Figure 3.**
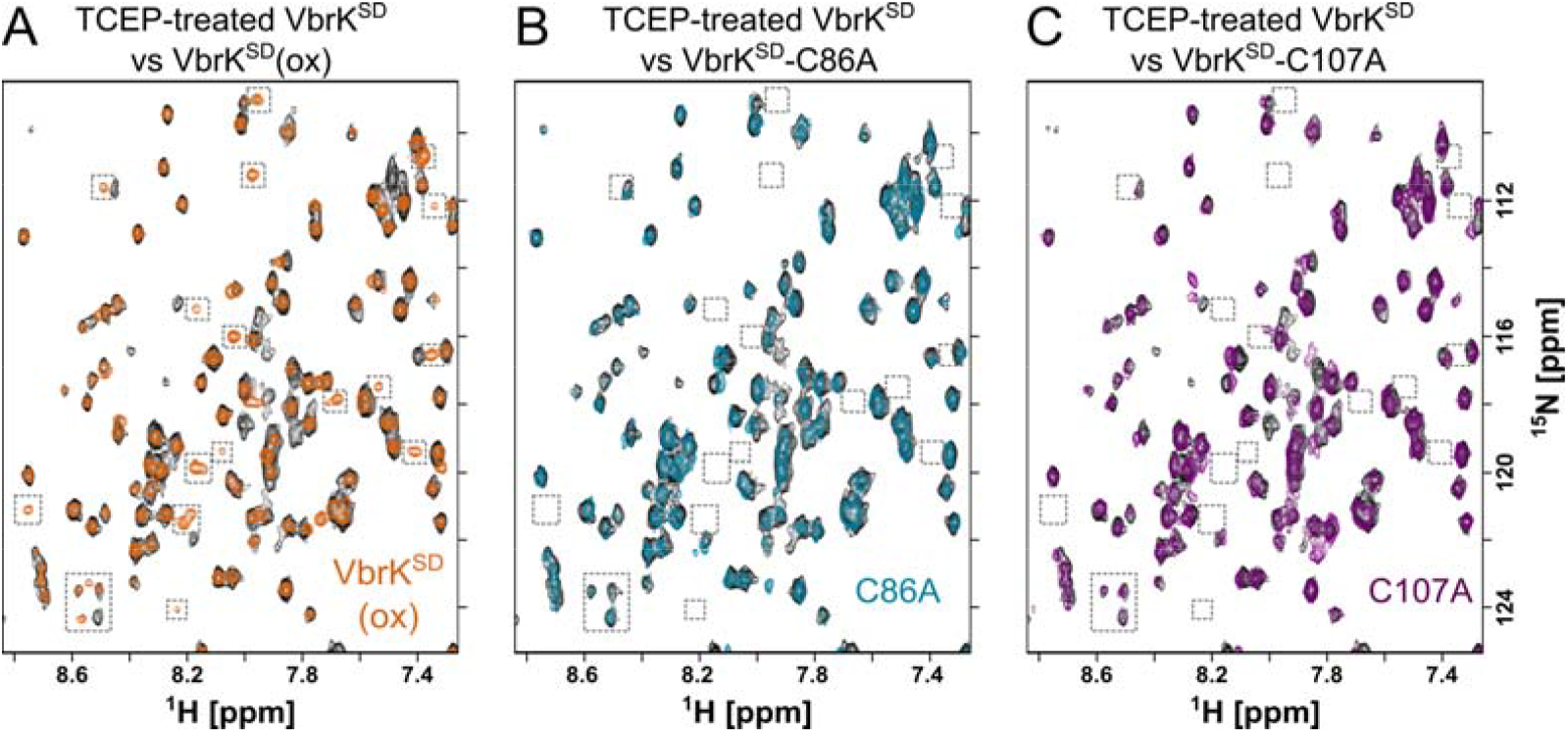
The ^1^H-^15^N HSQC spectra of wild-type VbrK^SD^, C86A and C107A mutants revealed the reduction of the C86-C107 disulfide bond upon TCEP-treatment. **A**. Overlay of ^1^H-^15^N HSQC spectra corresponding to untreated (ox for oxidized; orange) and TCEP-treated (black) wild-type (WT) VbrK^SD^. **B**. Overlay of ^1^H-^15^N HSQC spectra corresponding to TCEP-treated WT (black) and C86A (blue) VbrK^SD^. **C**. Overlay of ^1^H-^15^N HSQC spectra corresponding to TCEP-treated WT (black) and C107A (purple) VbrK^SD^ . Full spectra are shown in Figure S5.

We next tried to obtain a crystal structure of the reduced, antibiotic-bound form of VbrK^SD^. Co-crystallization experiments were not successful, but we used untreated protein crystals for soaking experiments, adding TCEP and penicillin G. We obtained the structure of the reduced VbrK^SD^ at 2.8 Å resolution (**Figure 4**.A and Table S2; PDB code 28QS). No electron density was detected for the antibiotic, but we clearly saw the absence of the disulfide bond between C86 and C107 (**Figure 4**.A). This disulfide bridge is formed in all previous crystal structures published of VbrK^SD^ from *V. parahaemolyticus* (PDB IDs: 7CUS and 7CJR) and its homologs in *V. cholerae* and *V. rotiferanus* (PDB IDs: 4R7Q, 7KB9, 7KB7, 7KB3, 7LA6, 7F2H and 7F2G)^9–12^. On the other hand, TCEP treatment did not affect the disulfide bond formed between C226 and C234 (**Figure 4**.A). The reduced structure was aligned against PDB 7CUS^9^ through the proximal subdomain (**Figure 4**.B). Even in the absence of bound ligand, the alignment shows a change in the relative orientation of the distal subdomain with respect to the proximal subdomain between oxidized and reduced forms of the protein which, in the full-length protein, might be required for initiation of signal transduction through the transmembrane region.

**Figure 4.**
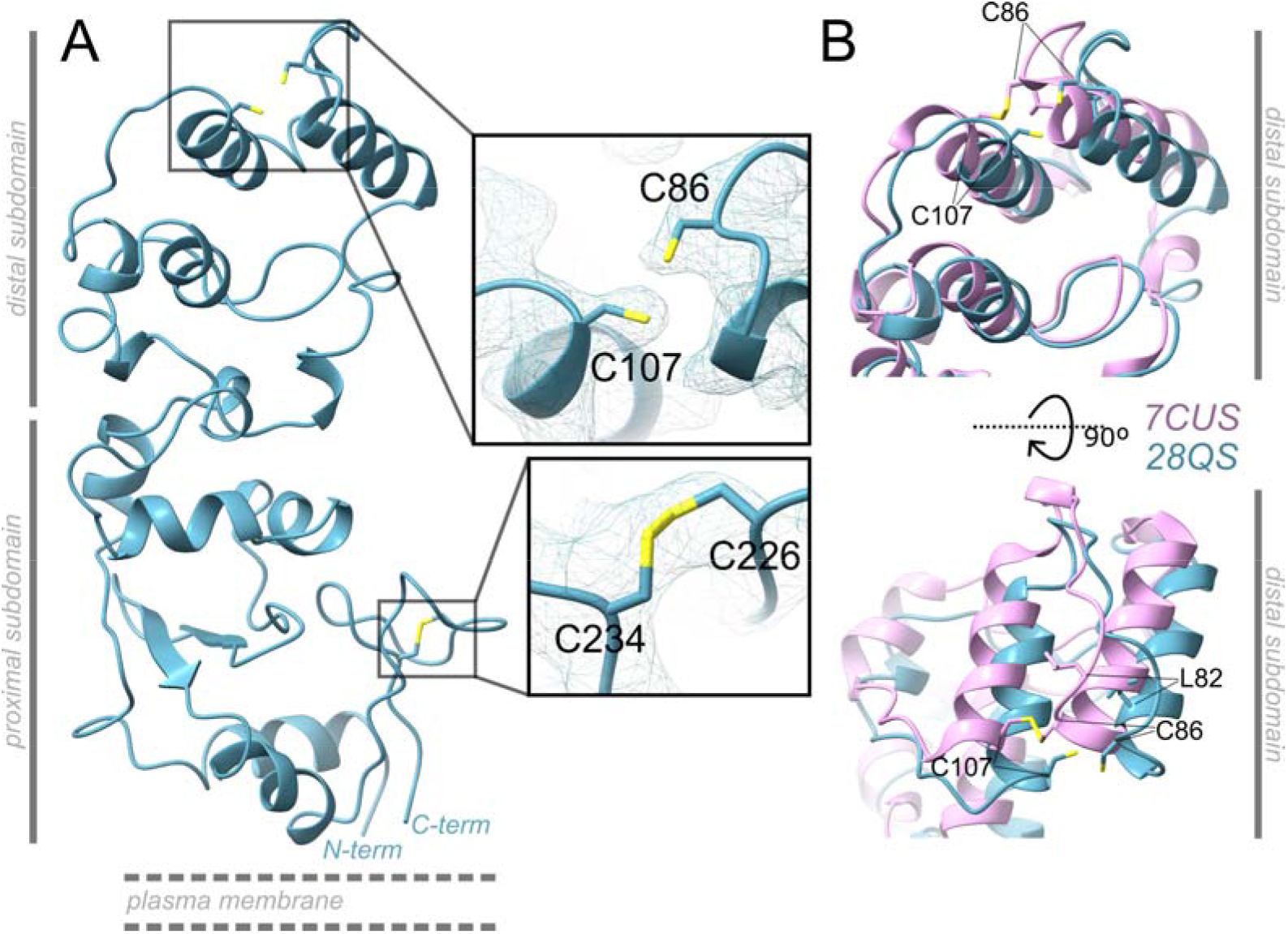
Crystal structure of reduced VbrK^SD^ showed the opening of the C86-C107 disulfide bridge. A. Crystal structure of VbrK^SD^ soaked with 3 mM TCEP and penicillin G, 2.8 Å resolution (PDB code 28QS). The electron density map of the regions containing C86, C107, C226 and C234 are shown in the insets. **B**. View of the distal domain of reduced VbrK^SD^ structure (28QS, blue) compared to the structure of oxidized VbrK^SD^ (7CUS, pink), when the α-carbons of the proximal domains are aligned.

The STD NMR experiments revealed that the interaction of β-lactams with VbrK^SD^ required reduction of the C86-C107 disulfide bond. Hence, STD NMR experiments were conducted to evaluate whether VbrK^SD^ C86A and C107A mutants still interacted with this class of antibiotics. We acquired STD NMR spectra of penicillin G in the presence of TCEP-treated wild-type VbrK ^SD^, C86A, C107A and C86A-C107A (**Figure 5**). All samples showed peaks at chemical shifts assigned to penicillin G protons. Hence, the mutations did not prevent the interaction of the antibiotic with VbrK^SD^. Interestingly, VbrK^SD^ C86A showed a higher perturbation of penicillin G proton signals, indicating an altered interaction mode. The C86A-C107A mutant showed the weaker STD NMR signals, comparable to the ones observed in assays with the L82A mutant, a previously reported mutation proposed to hinder the activation of the kinase by β-lactams^7^. According to the principles of STD NMR technique, lower intensity signals could be an indication of reduced affinity in a non-covalent interaction between C86A-C107A and L82A mutants and the ligand.

**Figure 5.**
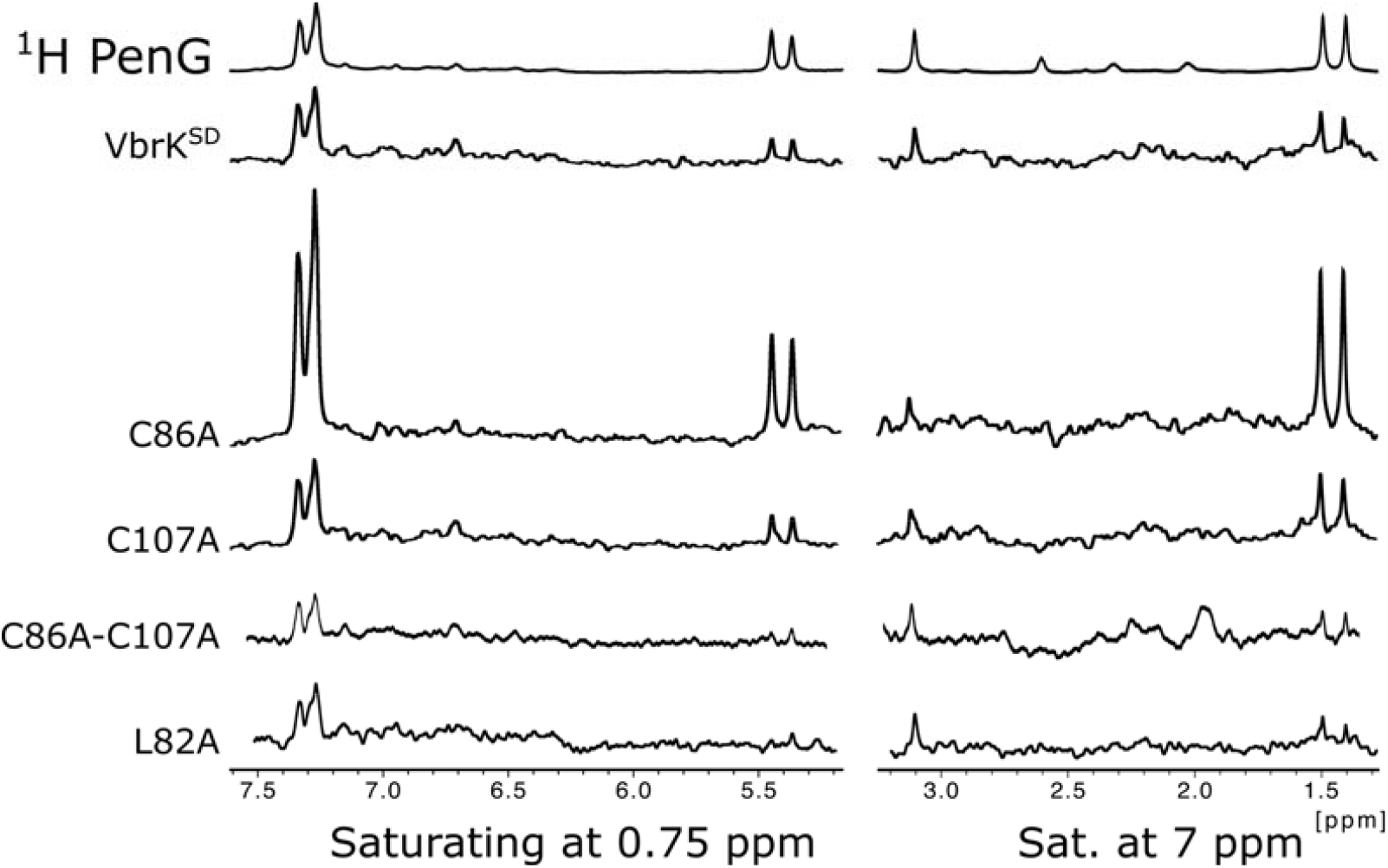
VbrK^SD^ C86A, C107A, C86A-C107A and L82A mutants form non-covalent complexes with penicillin G. STD-NMR experiments acquired on 400 µM penicillin G (PenG) in the presence of 20 µM TCEP-treated VbrK^SD^ WT, C86A, C107A, C86A-C107A and L82A. Top spectrum corresponds to the ^1^H NMR spectrum of 400 µM penicillin G; successive lower spectra correspond to STD-NMR spectra of penicillin G in the presence of VbrK^SD^ WT, C86A, C107A, C86A-C107A and L82A, saturating at 0.75 ppm (left) and 7 ppm (right).

### VbrK^SD^ forms a covalent adduct with β-lactams

Activation of a sensor protein requires formation of a stable ligand-sensor complex, one that is sufficiently long-lived to allow for signal transduction and phenotype manifestation. In the case of VbrK, S-nitrosylation of C86 activates the kinase VbrK, leading to its autophosphorylation, subsequent phosphotransfer to VbrR, and downstream gene regulation^8^. Hence, we wondered whether the non-covalent interaction of the sensor domain of VbrK with β-lactam antibiotics, that was observed in STD NMR experiments with reduced VbrK^SD^, could lead to formation of a covalent adduct between the ligand and the sensor protein. β-lactams inhibit transpeptidases through formation of stable covalent complexes, through a mechanism that involves a serine or cysteine residue as the nucleophile^17–20^. In the case of the β-lactam sensor proteins BlaR1 and MecR1, signal transduction is initiated by the nucleophilic attack of a serine residue on the carbonyl carbon of the β-lactam ring to form a stable-acylated species^21–25^. Given the need for disulfide bond reduction for interaction of VbrK^SD^ with β-lactam antibiotics, we reasoned that an acylation reaction could be involved. To test this hypothesis we incubated the protein with the fluorescent penicillin derivative Bocillin-FL^26^. Formation of a covalent VbrK^SD^-Bocillin-FL fluorescent adduct was evaluated by electrophoresis in denaturing polyacrylamide gels. Oxidized VbrK^SD^ did not react with Bocillin-FL, while TCEP-treated VbrK^SD^ formed a stable Bocillin-FL-acylated adduct that was detected as a fluorescent band in the gel (**Figure 6**.A). Then, we evaluated whether formation of the covalent adduct with TCEP-treated VbrK^SD^ was dependent on the presence of free cysteine residues. Reaction of the cysteines in TCEP-treated VbrK^SD^ with IAA or the cysteine labeling dye 7-Diethylamino-3-[4’-(iodoacetamido)phenyl]-4-methylcoumarin (DCIA) prior to incubation with Bocillin-FL impeded formation of the fluorescent β-lactam adducts (**Figure 6**.B). Proper reaction of DCIA with the reduced cysteine residues in VbrK^SD^ was confirmed by detection of the fluorescent DCIA-VbrK^SD^ adduct upon excitation with UV-light (**Figure 6**.B). Finally, we performed the same experiment in parallel with TCEP-treated wild-type VbrK^SD^, C86A, C107A, C86A-C107A and L82A (**Figure 6**.C). The C107A, C86A and L82A mutants showed lower accumulation of the Bocillin-FL-protein adduct than the wild-type protein. The C86A-C107A did not form a stable adduct with Bocillin-FL. Based on characterized mechanisms of β-lactam acylation of transpeptidases, a cysteine residue is a plausible nucleophile^18^. The absence of Bocillin-FL labelling when cysteines were blocked by IAA and DCIA and of the C86A-C107A VbrK^SD^ variant strongly suggested the involvement of a cysteine residue (C86 and/or C107A) in β-lactam acylation. On the other hand, the lower accumulation of Bocillin-FL-protein adduct for the L82A mutant demonstrated the importance of this residue for the stabilization of the covalent adduct (**Figure 6**.C).

**Figure 6.**
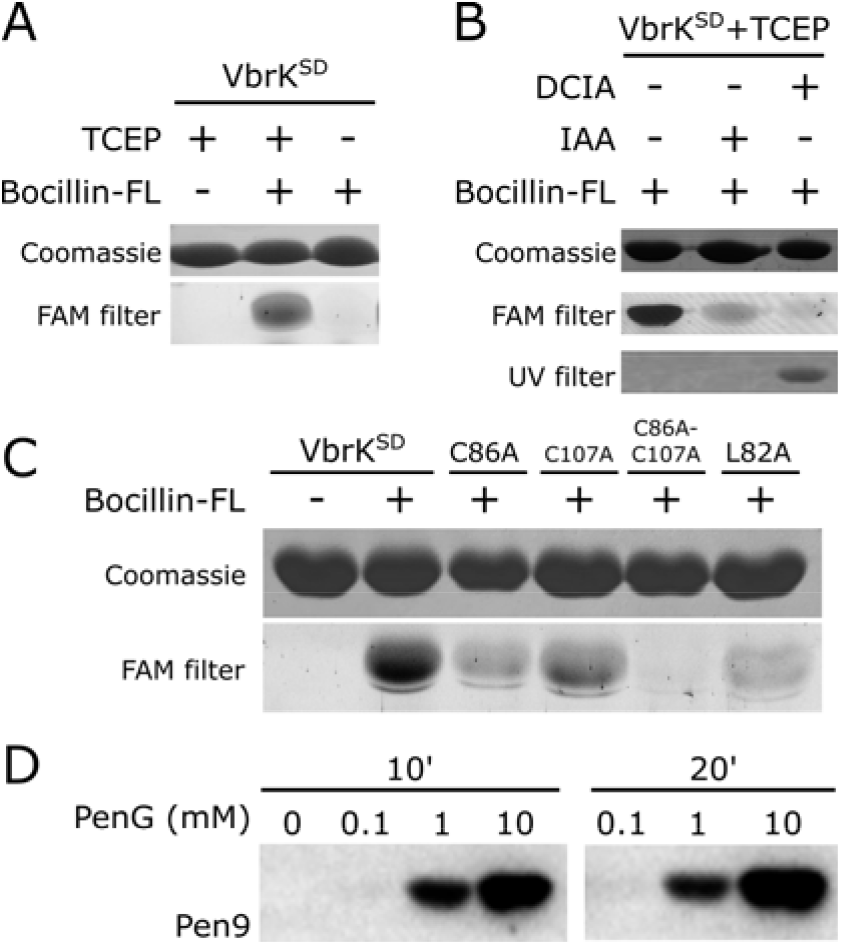
VbrK^SD^ forms a covalent complex with Bocillin-FL and Penicillin G. **A**. Non-TCEP treated and TCEP-treated VbrK^SD^ (100 µM) was incubated in the absence and in the presence of 100 µM Bocillin-FL for 20 min at 25 °C. **B**. TCEP-treated VbrK^SD^ (100 µM) was labeled at cysteine residues with IAA or DCIA, and then treated with 100 µM Bocillin-FL, for 20 min at 25 °C. **C**. TCEP-treated VbrK^SD^ WT, C86A, C107A, C86A-C107A and L82A (100 µM) were incubated with 100 µM Bocillin-FL, for 20 min at 25 °C. In A-C, proteins were resolved in 16% SDS-PAGE gels. Proteins bound to the fluorescent antibiotic Bocillin-FL were visualized using a FAM filter. In B, DCIA-bound proteins were visualized in a UV-transilluminator. The same gels were afterwards stained with Coomassie brilliant blue. **D**. Anti-Penicillin Western blot showing Penicillin G-labelled VbrK^SD^. TCEP-treated VbrK^SD^ (100 µM) was incubated in the absence and in the presence of 100 µM, 1 mM and 10 mM Penicillin G for 10 and 20 min at 25 °C. Full gel images are shown in Figure S8.

Formation of a covalent β-lactam-VbrK^SD^ adduct was also evidenced by Western blot upon incubation of TCEP-treated wild-type VbrK^SD^ with Penicillin G (PenG), where PenG-bound protein was detected through the recognition of its thiazolidine ring by the primary antibody Pen9 (**Figure 6**.D). Intact protein mass spectrometry analysis confirmed that VbrK^SD^ was acylated by Penicillin G, resulting in the accumulation of both singly and doubly PenG-labeled species (**Figure 7**). Covalent binding of one PenG molecule was observed at lower antibiotic concentrations and shorter incubation time. The amount of mono-acylated VbrK^SD^ increased with antibiotic concentration and incubation time. In these samples a second species was also observed, the di-acylated species. Top-down mass spectrometry analysis demonstrated that PenG acylated VbrK^SD^ on both C86 and C107 (Figure S9), confirming the nucleophilic attack of a cysteine residue (C86A and C107A) on the carbonyl carbon of the β-lactam ring, to form stable-acylated species. The mono-acylated VbrK^SD^ fraction consisted of two distinct species, with PenG attached to either C86 or C107, while the di-acylated fraction exhibited modification at both cysteine residues. PenG acylation on C86 could also be confirmed upon incubation of VbrK^SD^ C107A with PenG (Figure S10). No covalent adduct was detected with the oxidized wild-type VbrK^SD^ sample (Figure S11) nor with the VbrK^SD^ C86A-C107A mutant (Figure S12).

**Figure 7.**
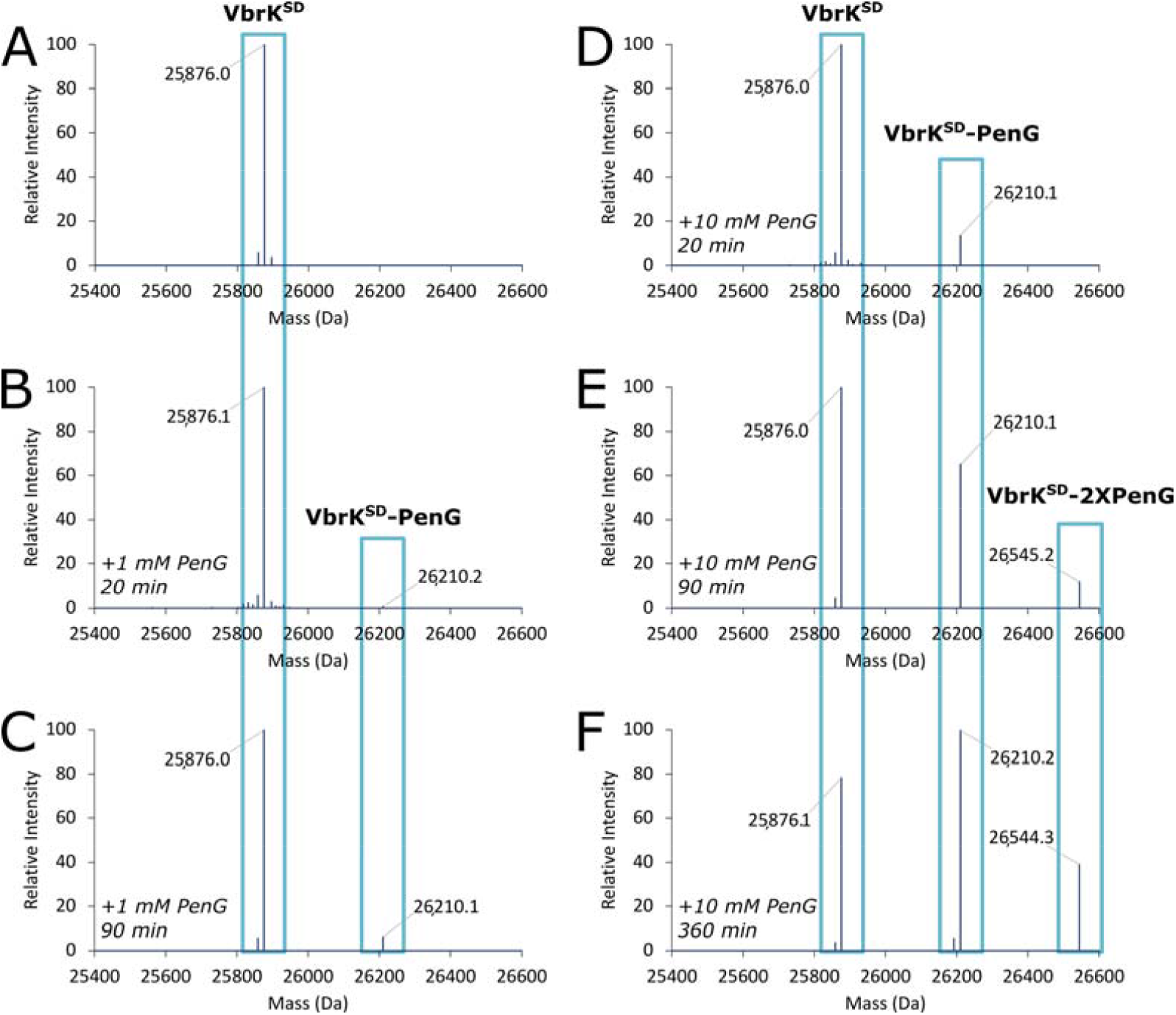
Intact protein mass spectrometry analysis confirms that VbrK^SD^ forms a covalent complex with Penicillin G, by acylation of C86 and C107. **A**. Deconvolved mass spectrum of TCEP-treated VbrK^SD^ (100 µM). **B-C**. Deconvolved mass spectrum of TCEP-treated VbrK^SD^ (100 µM) incubated with 1 mM Penicillin G for 20 min (B) and 90 min (C), at 25 °C. **D-F**. Deconvolved mass spectrum of TCEP-treated VbrK^SD^ (100 µM) incubated with 10 mM Penicillin G for 20 min (D), 90 min (E) and 6 h (F). A-F display the monoisotopic mass values in Da with mass accuracy of 5 ppm.

### VbrK^SD^ slowly hydrolyzes β-lactams, following a transpeptidase-like mechanism

VbrK is activated (undergoes autophosphorylation) in the presence of β-lactams^7^. If covalent modification of the sensor domain of VbrK were the first step in the signal transduction mechanism, and the β-lactam–acylated protein represented the active kinase conformation, then de-acylation would allow VbrK to return to its inactive state and the system to shut down.

To determine if VbrK^SD^ is recycled by de-acylation, we followed the hydrolysis of the chromogenic cephalosporin nitrocefin using a stopped-flow spectrophotometer equipped with a photodiode array detector (**Figure 8**). The electronic spectrum of nitrocefin presents a maximum of absorbance at 392 nm, while hydrolyzed nitrocefin presents a band with maximum at 495-500 nm. Hydrolysis was negligible in the presence of nanomolar concentrations of TCEP-treated VbrK^SD^ (not shown). However, incubation of nitrocefin with higher concentrations of TCEP-treated VbrK^SD^ (micromolar protein concentrations) resulted in a clear decrease in the intensity of absorbance at 392 nm over time, with a concomitant increase in the absorbance at 495 nm, which demonstrated that VbrK^SD^ catalyzed the hydrolysis of nitrocefin (**Figure 8**.B). The observed low hydrolytic activity, contrasted with that of efficient β-lactamases^27^ but was consistent with the turnover rate of β-lactam activated sensor proteins^21,25^. Oxidized VbrK^SD^ did not hydrolyze nitrocefin (**Figure 8**.A). We then tested nitrocefin hydrolysis by the C86A and C107A VbrK^SD^ mutants. VbrK^SD^ C86A did not present β-lactamase activity, while the activity of the C107A mutant was similar to the activity of the wild-type sensor domain (**Figure 8**.C and D).

**Figure 8.**
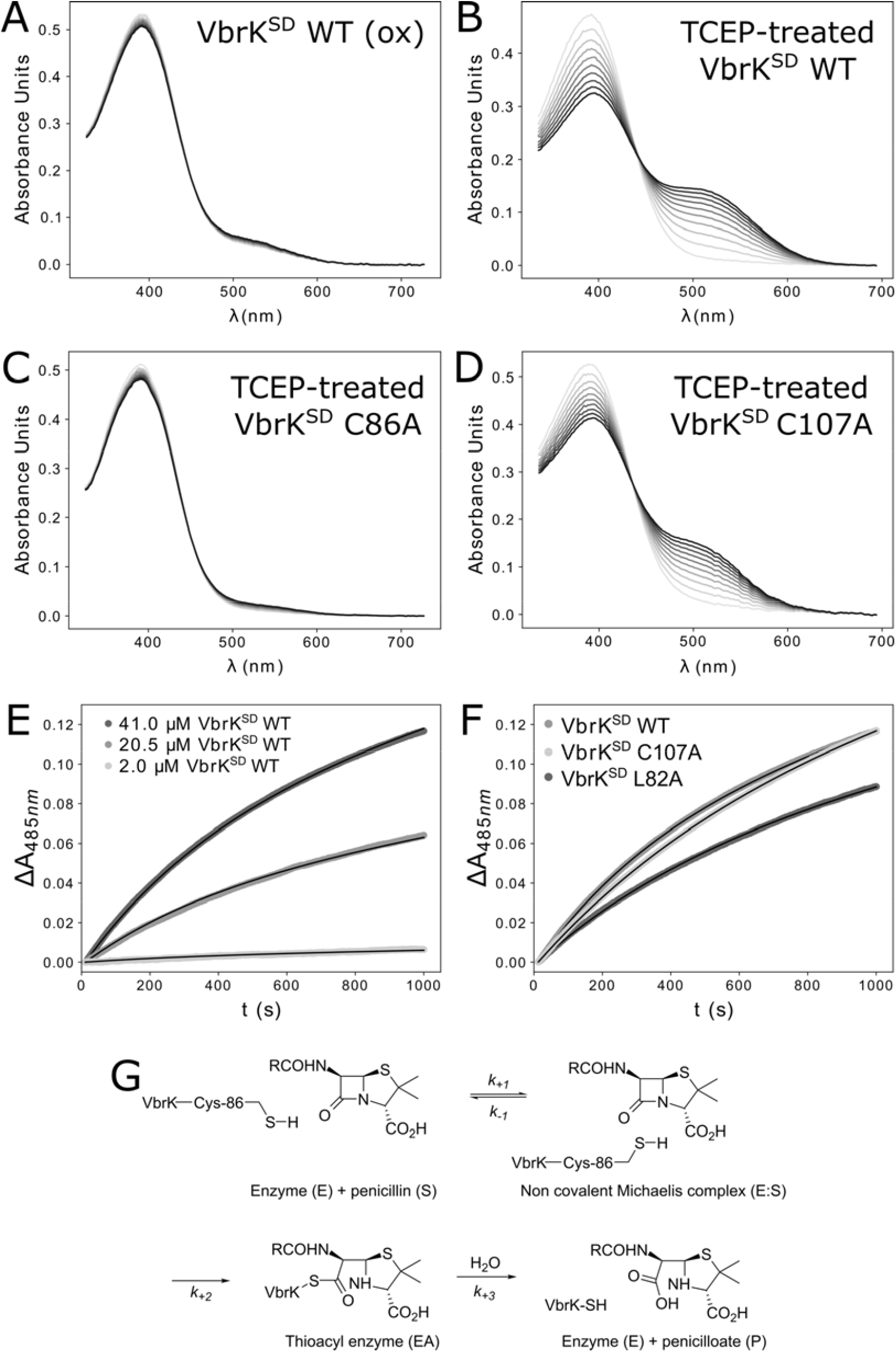
VbrK^SD^ hydrolyzed the cephalosporin nitrocefin. **A**. UV-visible absorbance spectra over 1000 s (grey to black, every 100 s) of 41 µM nitrocefin incubated with 41 µM oxidized VbrK^SD^. B. - D. Same experiment as in A but with TCEP-treated VbrK^SD^, C86A and C107A, respectively. E. Change in Absorbance at 485 nm (ΔA_485 nm_ = A_485 nm,t_ – A_485 nm,2 ms_) as a function of time, for the reaction of 41 µM nitrocefin with three different concentrations of TCEP-treated VbrK^SD^ WT (2, 20.5 and 41 µM); black lines correspond to the calculated curve using kinetic parameters derived from differential equation analysis with the model described in G. k_+1_ was fixed to the diffusional limited value of 1 x 10^8^ s^-1^ and the differential extinction coefficient for hydrolyzed free nitrocefin at 485 nm was 17,420 M^-1^ cm^-1 28^. To obtain a good fit of the data, the extinction coefficient for nitrocefin covalently bound to the enzyme was also fit. Estimated kinetic parameters were: k_−1_ = (50 ± 130) x 10^3^ s^-1^; k _2_= (1.0 ± 0.3) x 10^-3^ s^-1^; k_3_= (2.50 ± 0.02) x 10^-3^ s^-1^; Δℰ_EA_ = (70 ± 1) x 10^3^ M^-1^ cm^-1^. Differences between the molar absorptivity of free nitrocefin and protein-bound nitrocefin was previously reported^18^. F. Change in Absorbance at 485 nm as a function of time, for the reaction of 41 µM nitrocefin with 41 µM TCEP-treated VbrK^SD^ WT, C107A and L82A variants; kinetic parameters shown in Table S3. G. Proposed mechanism of β-lactam hydrolysis catalyzed by TCEP-treated VbrK^SD^. The scheme is presented for a penicillin, but the time courses of hydrolysis of the cephalosporin nitrocefin fit to the same mechanism, as shown in F.

The progress curves of nitrocefin hydrolysis catalyzed by three different concentrations of TCEP-treated VbrK^SD^ (2-41 µM; **Figure 8**.E) were fit to different models. The best fit was obtained with a system of differential equations representing the reaction mechanism depicted in **Figure 8**.G. In this model, VbrK^SD^ initially forms a non-covalent Michaelis complex with the substrate (E:S), with subsequent formation of a covalent antibiotic-enzyme complex (EA). The covalent intermediate, then slowly releases the hydrolyzed antibiotic (P) and regenerates the free enzyme (E). The fitted kinetic parameters are presented in the legend of **Figure 8** and in Table S3. The time-courses of hydrolysis of nitrocefin by the C107A mutant could be fit to the same model as the nitrocefin hydrolysis data catalyzed by wild-type VbrK^SD^ (**Figure 8**.F and Table S3). The neglectable hydrolytic activity of C86A mutant evidenced this residue is essential for hydrolysis of nitrocefin, suggesting it might be essential for the complete signaling cycle to occur. The L,D-transpeptidase Ldt_fm_, from *Enterococcus faecium*, forms a covalent adduct with β-lactams through a mechanism involving nucleophilic attack by the thiol of a cysteine residue to form a thioester bond, yielding a cysteine-acylated intermediate^18^. Release of the hydrolyzed β-lactam from VbrK^SD^ -de-acylation of the covalent β-lactam-sensor adduct - would allow for regeneration of the pool of free sensor protein, thereby enabling reset of the two-component system once the β-lactam signal is no longer present.

It was previously shown that VbrK mutant L82A does not auto-phosphorylate upon treatment with the β-lactam carbenicillin and that this mutation abolishes the ability to produce β-lactamase by *Vibrio parahaemolyticus*^*7*^. Based on these results, it was proposed that L82 is critical for β-lactam recognition^7^. In this study, STD NMR experiments showed interaction of β-lactams with VbrK^SD^ L82A, only after TCEP-treatment, as observed with the wild-type sensor domain (**Figure 5**). The intensity of the STD signal was lower than for the wild-type domain, indicating the non-covalent interaction still occurred but was affected. In addition, VbrK^SD^ L82A catalyzed nitrocefin hydrolysis with kinetic parameters similar to those of the wild-type sensor domain (**Figure 8**.F and Table S3). Hence, VbrK^SD^ L82A formed a non-covalent complex with β-lactams (detected in the STD NMR experiments) and hydrolyzed them, but it accumulated a lower amount of the covalent adduct with Bocillin-FL (**Figure 6**.C and Figure S13). These results suggest that L82 is required for stabilization of the covalent adduct. Together with the fact that the L82A substitution impairs β-lactam induction of auto-phosphorylation/phosphotransfer activities in VbrK^7^, we propose that the β-lactam–acylated sensor protein is the active form of the kinase, required for activation of the VbrK/VbrR system.

### Full-length VbrK reacts and forms a covalent adduct with the β-lactam Bocillin-FL

We have documented that the sensor domain of VbrK formed a covalent adduct with the β-lactam Bocillin-FL upon reduction of the C86-C107 disulfide bond by incubation of purified VbrK^SD^ with TCEP. If activation of the kinase VbrK by β-lactams were due to formation of a stable thioester following nucleophilic attack by C86 and C107 on the carbonyl carbon of the β-lactam ring, then the C86-C107 disulfide bond would be expected to be reduced in the periplasm. To test this hypothesis, full-length VbrK was expressed in *E. coli* and the reaction with Bocillin-FL was evaluated upon incubation of whole cells, spheroplasts and membrane preparations with this antibiotic, which were then analyzed by SDS-PAGE and fluorescence scanning. The Bocillin-FL signal of the samples produced with whole cells was low, presumably due to a poor amount of Bocillin-FL crossing the outer membrane (data not shown). An intense band of Bocillin-FL-labeled protein was observed at the position in the gel expected for VbrK (**Figure 9**; VbrK mass was *ca*. 54 kDa after cleavage of the signal peptide, Figure S2). This band was detected on spheroplasts and membrane preparations from *E. coli* cells expressing full-length VbrK (**Figure 9**). This result showed that the periplasmic sensor domain of full-length VbrK was in a redox state that allowed for the reaction with β-lactam antibiotics to occur, producing the accumulation of the stable acylated VbrK species. The amount of fluorescently-labelled VbrK increased when the preparation of membrane proteins was incubated with TCEP prior to incubation with Bocillin-FL. Hence, overexpression of VbrK in *E. coli* membranes resulted in a mixed protein population: a fraction of VbrK that directly reacted with the antibiotic, and a second population that was labelled only after incubation with a reducing agent. This could be due to expression levels of VbrK surpassing the capacity of periplasmic disulfide oxidoreductases to reduce the C86-C107 disulfide bond in VbrK, as we discuss below. The identity of the fluorescent band attributed to VbrK was confirmed by Western blot with a His-Tag-specific antibody (Figure S14).

**Figure 9.**
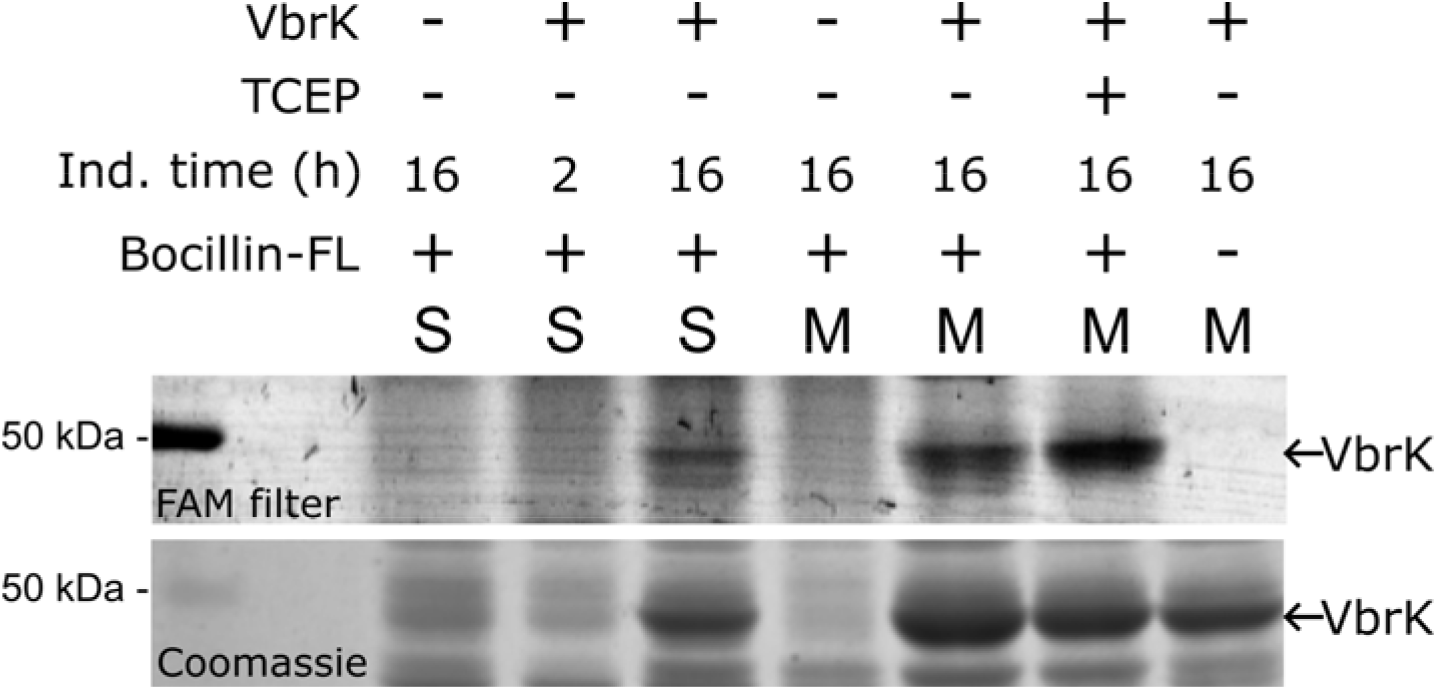
Intrinsic reduction of VbrK in cellular membrane contexts enables covalent Bocillin-FL labeling. Full-length VbrK was labeled with Bocillin-FL in spheroplasts (S), and membrane protein extracts (M), followed by analysis on a 16% SDS-PAGE gel. Expression of VbrK in E. coli BL21 Star™ DE3 was induced with 10 µM IPTG during 2 or 16 h, as indicated. Samples were incubated with 80 μM Bocillin-FL (final concentration) for 20 min at 25°C. Spheroplasts were not incubated with any reducing agent. Membranes were labelled with Bocillin-FL as isolated and after treatment with 3 mM TCEP for 10 min at 25 °C. Bocillin-FL-labelled proteins were detected using a FAM filter (top image). Bottom image: same gel stained with Coomassie Brilliant Blue, as a loading control. VbrK (ca. 54 kDa) is pointed with an arrow. Full gel images and a Western blot with α-His-Tag antibody are shown in Figure S14.

## Discussion

In this work, we characterized the interaction of β-lactam antibiotics with VbrK, a histidine kinase from *V. parahaemolyticus* that is activated in the presence of these antibiotics and that initiates a signaling cascade that results in expression of the serine β-lactamase CARB^6,7^. We documented that β-lactams bind to VbrK expressed in membranes, and more specifically, that the DUF3404 (VbrK^SD^) sensor domain is a novel class of redox-gated β-lactam recognition module.

VbrK has a signal peptide which we show is cleaved upon translocation to the inner membrane, resulting in an N-terminal sensor domain, VbrK^SD^, connected through a transmembrane *α*-helix to the cytoplasmic kinase/phosphatase domain. VbrK^SD^ has two disulfide bonds, one located in the subdomain that would be proximal to the membrane and one located in the membrane-distal subdomain (the C86-C107 bond; **Figure 10**). STD NMR experiments demonstrated that reduction of one disulfide bond is required for non-covalent β-lactam binding to VbrK^SD^, indicating that the binding site is not available in the oxidized sensor domain, consistent with a redox-gated, induced-fit mechanism of ligand recognition. This finding reconciles previous conflicting reports, which can now be attributed to differences in protein redox state during preparation^7,10^. The structure of reduced VbrK^SD^ in combination with ^1^H-^15^N HSQC and mass spectrometry experiments on wild-type, C86A and C107A mutant versions provide evidence that the C86-C107 disulfide bond is the one reduced.

**Figure 10.**
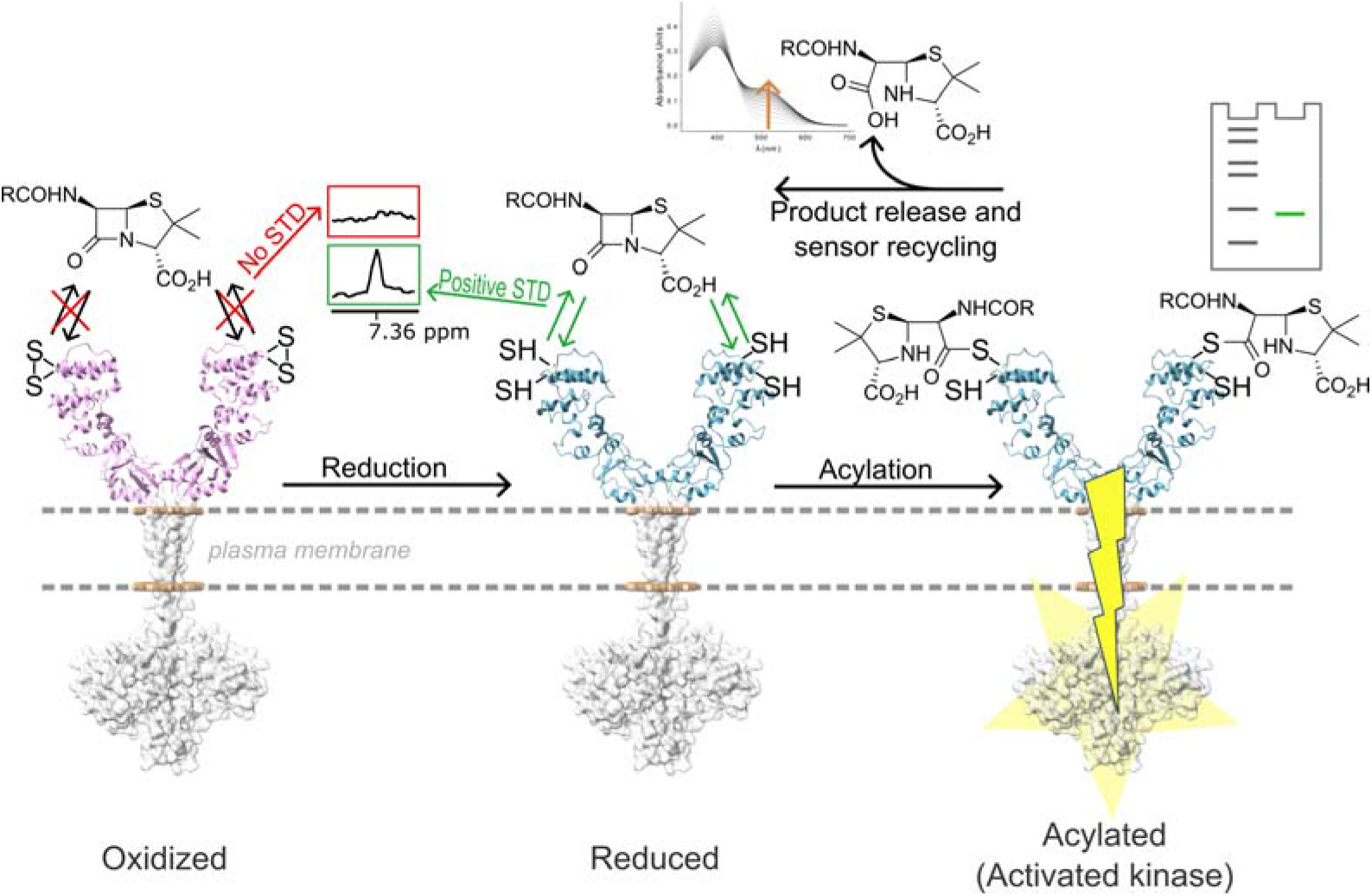
Proposed mechanism of recognition of β-lactam antibiotics by VbrK. Pink oxidized VbrK^SD^ structure corresponds to PDB 7CUS, blue reduced VbrK^SD^ structure corresponds to PDB 28QS (this study). Transmembrane and cytoplasmic histidine-kinase domains is an AlphaFold2 model of VbrK used only for representation purposes. In the oxidized species, we show schematically the C86-C107 disulfide bond, in the membrane-distal subdomain of VbrK^SD^; this is the disulfide bond that is reduced for the interaction with β-lactam antibiotics to occur.

Combining assays with a fluorescent penicillin derivative, penicillin-specific antibodies, stopped-flow kinetics, mutagenesis and mass spectrometry, we demonstrated that initial non-covalent binding of the antibiotic to VbrK^SD^ leads to a covalent modification by β-lactams. Acylation by β-lactam antibiotics was observed on C86 and, to a lower extent, on C107. A conserved strategy for the formation of covalent β-lactam-protein adducts involves the nucleophilic attack on the β-lactam carbonyl carbon by an active-site serine residue, in PBPs and serine β-lactamases^29–31^ or by a cysteine, in L,D-transpeptidases^18^, resulting in β-lactam ring scission and the formation of a stable acyl–enzyme adduct. The evidences of β-lactam labelling of VbrK on C86 and C107 support a mechanism analogous to that in L,D-transpeptidases, in which the thiol group of a cysteine residue attacks the β-lactam carbonyl, forming a thioester intermediate. Stopped-flow kinetic studies demonstrated that the reaction with the β-lactam antibiotic nitrocefin involves accumulation of an acyl-enzyme adduct and slow release of the hydrolyzed antibiotic to regenerate the free protein.

Kinetic studies on mutant versions of VbrK^SD^ unveiled differences in the roles of C86 and C107. Hydrolysis of nitrocefin by VbrK^SD^ C86A is negligible while wild-type like β-lactam hydrolysis is evidenced in the presence of C86 (C107A mutant). The L82A mutation does not impair β-lactam binding or hydrolysis but prevents accumulation of the covalent adduct. Hence, we propose that the previously reported inability of the L82A mutant to activate kinase signaling^7^ is due to its deficiency in stabilizing the β-lactam-acylated adduct. This highlights the importance of the acylated intermediate in VbrK activation. We propose that β-lactam acylation leads to the kinase-active state. Two-component systems should remain active while the signal is present. In the case of the VbrK/VbrR system, its β-lactam-induced activation leads to CARB expression. This β-lactamase will eventually hydrolyze the β-lactam antibiotic, eliminating the signal that activates VbrK/VbrR. However, the system will only shut down when VbrK is no longer in its kinase-active conformation, the β-lactam acylated species. Hence, VbrK inactivation would require release of the hydrolyzed antibiotic, that is de-acylation of the β-lactam-VbrK adduct (**Figure 10**). The nitrocefin assays allowed us to document that hydrolysis eventually occurs, at a rate consistent with that expected for β-lactam sensor proteins^21,25^, enabling the system to reset once the signal is removed. The differences observed in hydrolysis between VbrK^SD^ C86 and VbrK^SD^ C107 suggest that C86 is essential for the hole signaling cycle to take place.

VbrK is present in all *Vibrio* species, with 60–90% sequence identity among them. VxrA from *V. cholerae* is a VbrK homolog, with 64.1% sequence identity in their sensor domains^12^. As is the case for VbrK, VxrA positively autoregulates its expression and contributes to enhanced survival in response to β-lactam exposure^32^. The structures of VxrA^SD^ (PDB IDs: 7KB7, 7KB9, 7KB3 and 4R7Q) show two disulfide bonds, equivalent to the ones formed in VbrK^SD^, with the C101-C122 disulfide bond located in the membrane-distal subdomain, equivalent to the C86-C107 disulfide bond in VbrK. The mutation C101A-C122A in VxrA in *V. cholerae* resulted in reduced *vxrA* expression and significant reduction in survival following penicillin G treatment. These results agree with the role here proposed for cysteines 86 and 107 in VbrK.

Recombinant full-length VbrK efficiently reacted with Bocillin-FL both in spheroplasts and membranes, indicating that the C86-C107 disulfide bond is reduced in the periplasm of *E. coli*. Given that the periplasm is an oxidizing environment, the cysteine residues of periplasmic proteins are frequently involved in disulfide bonds. It is then likely that the two disulfide bonds are formed in the sensor domain of VbrK. However, disulfide bonds present variable stabilities, with structural disulfide bonds being more stable than disulfide bonds formed between cysteine residues involved in catalysis^33^. One possibility is that the redox potential of the C86-C107 disulfide bond allows for the reaction of C86 and C107 with β-lactams (as reported here) or with other signaling molecules (*e*.*g*. nitrite^8^) to occur. Another possibility is that disulfide bond formation and reduction in VbrK is regulated by disulfide oxidoreductases, as is the case for other periplasmic proteins and domains^33,34^. In the light that the interaction of β-lactams with VbrK is dependent on reduction of the C86-C107 disulfide bond in its sensor domain, we can envision two new alternatives to restore the efficacy of β-lactams in *Vibrio*: impeding reduction of the C86-C107 disulfide bond or impeding formation of the β-lactam-VbrK adduct.

## Conclusion

We identify the VbrK sensor domain as a previously unrecognized β-lactam–reactive module whose activity is governed by cysteine redox state. Reduction of the C86-C107 disulfide bond in the periplasmic sensor domain of VbrK is a prerequisite for non-covalent β-Lactam binding, which then results in covalent modification at C86 and C107. Mutagenesis studies indicate that β-Lactam acylation stabilizes the kinase-active conformation, whereas de-acylation would enable system shutdown. Together, these results define a new mechanism of histidine kinase regulation and open opportunities for targeting β-lactam sensing pathways to combat antibiotic resistance in *Vibrio*. New strategies to restore β-lactam efficacy in *Vibrio* can be envisioned, including inhibition of adduct formation or targeting redox regulation that control VbrK activation.

## Materials and methods

### Reagents

Lisogeny Broth (LB) and LB-agar were from BD Biosciences. Plasmid preparations were carried out using *EasyPure*^*®*^ Plasmid Miniprep Kit (TransGen Biotech). DNA fragments were purified using *EasyPure*^®^ *Quick Gel Extraction Kit* (TransGen Biotech). Reagents used for buffers were from Sigma, Merck, Thermo Fisher Scientific. Restriction enzymes were from New England Biolabs and Promega; T4 DNA ligase was from TransGen Biotech. PCR reactions for cloning were done with Phusion Hot Start II DNA Polymerase (Thermo Fisher Scientific).

### Cloning of VbrK from V. parahaemolyticus and its variants

The synthetic gene encoding full length VbrK protein from *V. parahaemolyticus* (UniProt accession number Q87HP3) was obtained from GenScript (United States). Its sequence was optimized for expression in *E. coli* and flanked by NdeI and XhoI restriction sites. The synthetic gene was cloned into pET24a(+) after NdeI/XhoI digestion in phase with a C-terminal H6X tag, to yield plasmid pET24a(+)::*vbrK*. VbrK^SD^ constructs spanning residues 2-240 and 25-240 were amplified by PCR using pET24a(+)::*vbrK* as a template and oligos including NcoI and EcoRI restriction sites at the 5’ and 3’ ends respectively. The fragments were cloned into a, NcoI/EcoRI digested, modified pET28a(+) vector, which allows the generation of fusion proteins with a histidine hexapeptide at its C-terminal end, preceded by a thrombin protease cleavage site (pET28a(+), here after). The plasmids obtained were pET28a(+)::*vbrK*^SD^*(2-240)* and pET28a(+)::*vbrK*^SD^*(25-240)*. Later during this project, the thrombin digestion site on pET28a(+)::*vbrK*^SD^*(2-240)* was replaced by a TEV protease digestion site by PCR directed mutagenesis, to give plasmid pET28a(+)::*vbrK*^SD^*(2-240)_TEV-H6x*. Point mutants were generated by site-directed mutagenesis using pET28a(+)::*vbrK*^SD^*(2-240)_TEV-H6x* as the template. All sequences were confirmed by DNA sequencing service of Macrogen (Seoul, Republic of Korea). DNA oligo sequences are listed in Table S1.

### Expression of VbrK and preparation of VbrK-containing membrane protein extracts

Wild-type VbrK was expressed in freshly transformed *E. coli* BL21 Star™ (DE3) cells harboring the pET24a(+)::*vbrK* plasmid, encoding VbrK with a C-terminal H6X tag. A 1000-mL LB culture in a 5 L Erlenmeyer flask was inoculated at 1:50 dilution from an overnight saturated culture and supplemented with 50 µg/mL kanamycin. Cultures were grown at 37°C with shaking (180 rpm) until OD_600_ = 0.4, then cooled to 20°C and grown to OD_600_ = 0.6. Expression was induced with 10 µM or 500 µM IPTG, as indicated, and cultures were incubated for 2, 4 and 16 h at 20 °C with shaking (180 rpm).

Cells were harvested by centrifugation at 4700 *g*, for 20 min at 4 °C, resuspended in 40 mL lysis buffer (12 mM Na_2_HPO_4_ pH 7.4, 500 mM NaCl, 2.7 mM KCl, 0.5 mM PMSF, 0.1 mM EDTA, 10% (v/v) glycerol), and disrupted by three passes through an Emulsiflex C3 high-pressure homogenizer (10,000–15,000 psi). The lysate was then ultracentrifuged at 100,000 *g* for 1 h at 4 °C in an Optima™ L-90K ultracentrifuge (Beckman Coulter) using a Ti90 rotor. The soluble fraction was separated, and the membrane pellet was resuspended in 12 mM sodium phosphate buffer pH 7.4, 500 mM NaCl, 2.7 mM KCl, 20 mM imidazole, 10% (v/v) glycerol buffer using a Potter-Elvehjem PTFE pestle and glass homogenizer (Sigma-Aldrich), to obtain a suspension of membrane vesicles containing 300 mg/mL total membrane protein. Membranes were flash-frozen in liquid N_2_ and stored at −80 °C until use.

### VbrK spheroplasts preparation

All steps were performed at 4 °C. A 10-mL aliquot of induced culture expressing VbrK was centrifuged 4,700 *g* for 15 min at 4 °C, and the cell pellet was washed once with 1 mL of 50 mM Tris-HCl, pH 8.0, 150 mM NaCl. Cells were then pelleted at 4,700 *g* for 15 min at 4 °C and resuspended in 10 mL of Lysozyme Buffer (20 mM Tris-HCl, pH 8.0, 0.1 mM EDTA, 20% (w/v) sucrose, 0.5 mM PMSF, 1 mg/mL lysozyme) and incubated for 30 min at 4 °C. Spheroplasts were pelleted at 4,700 *g* for 15 min at 4 °C, resuspended in 10 mL of 20 mM Tris-HCl, pH 8.0, 20% (w/v) sucrose, centrifuged again at 4,700 *g* for 15 min at 4 °C, and finally resuspended in 1 mL of 20 mM Tris-HCl, pH 8.0, 10 mM CaCl_2_. Spheroplasts were treated right after preparation.

### Expression and purification of wild-type VbrK^SD^ and mutant versions

Wild-type and mutant versions of VbrK^SD^ were expressed in *E. coli* BL21 Star™ (DE3) cells freshly transformed with plasmids pET28a(+)::*VbrK*^SD^*(2-240)* coding for the wild-type version (with Thrombin site for assays shown in Figures S1.A; with TEV site for all subsequent assays, including Figure S1.B) or the corresponding mutant version (C86A, C107A, C86A-C107A, L82A; all with TEV site). From a saturated culture, a 1:50 dilution was made in 1000 mL of LB medium, in 5 L-Erlenmeyer flasks, supplemented with 50 µg/mL kanamycin. When ^15^N-labelled protein was produced, protein expression was performed on M9 medium (0.5 g/L NaCl, 6 g/L Na_2_HPO_4_.7H_2_O, 3 g/L KH_2_PO_4_ pH 7.4, 0.4% (p/v) glucose, 1 mM MgSO_4_, 0.33 mM CaCl_2_, 0.1% (p/v) NH_4_Cl) supplemented with ^15^NH_4_Cl as nitrogen source. The culture was incubated at 37 °C with shaking (200 rpm) until reaching an OD_600nm_= 0.4, when the temperature was lowered to 20 °C. The culture was further incubated at 20 °C until it reached an OD_600nm_= 0.6. At this point, protein expression was induced by the addition of 10 µM IPTG; protein expression was induced by incubation of the culture for 16 h at 20 °C with shaking (200 rpm). *E. coli* cells were harvested by centrifugation at 4700 *g*, for 20 min at 4 °C, resuspended in 20 mL of lysis buffer (50 mM Tris-HCl, pH 8, and 500 mM NaCl, 30 mM Imidazole, and 0,5 mM PMSF, 0,1 mM EDTA, 3 mM 2-mercaptoethanol and 5% (v/v) glycerol). Cells were disrupted by sonication: subjected to 10 pulses with 20% power in a 600-Watt Sonics GEX-600 sonicator (Sonics & Materials); each pulse was applied for 15 s, leaving 60 s intervals between pulses, keeping the solution in a water-ice bath). Cell debris were separated by centrifugation at 4 ºC, 21000 *g* for 20 minutes. The protein of interest was bound to 1 mL of Ni-Sepharose 6 Fast Flow resin (Cytiva) equilibrated with 50 mM Tris-HCl, pH 8, 500 mM NaCl, 30 mM imidazole, 0,5 mM PMSF, 3 mM 2-mercaptoethanol and 5% (v/v) glycerol. The resin was washed with fifteen column volumes of the same buffer and fifteen column volumes of binding buffer supplemented with 40 mM of imidazole. VbrK^SD^ was eluted with 10 column volumes of 50 mM Tris-HCl, pH 8, 500 mM NaCl, 3 mM 2-mercaptoethanol, 5% (v/v) glycerol and 170 mM imidazole. The presence of the protein of interest in the different fractions was checked by absorption at 280 nm and SDS-PAGE. *Tobacco Etch Virus nuclear-inclusion-a endopeptidase* (TEV) protease was added to the pooled fractions in a 1:50 molar ratio and the mixture was dialyzed versus 1 L of 20 mM Tris-HCl, pH 8, 300 mM NaCl, 3 mM 2-mercaptoethanol, and 5% (v/v) glycerol for 16 hours at 4°C using 14 kDa cut off dialysis tubing (Sigma-Aldrich). In the case of thrombin digested samples 0.1 mg of thrombin was added every 1 mg of purified protein and the mixture was dialyzed with the same protocol as for TEV digestion. After dialysis, the protein solution was loaded on Ni-Sepharose resin equilibrated in 20 mM Tris-HCl, pH 8, 300 mM NaCl, 3 mM 2-mercaptoethanol, 5% (v/v) glycerol. The protein of interest was collected on the flow-through and a fifteen-column-volumes wash with equilibration buffer. Flow-through and wash were pooled and concentrated up to 8 mg/ml. The protein was purified by a size exclusion chromatography step with a Superdex 200 Increase 10/300 GL column (Cytiva) equilibrated in 20 mM Tris-HCl pH 8, 150 mM NaCl. The presence of the protein of interest in the different fractions was detected by absorption at 280 nm and SDS-PAGE. The purest protein fractions were pooled. 1X cOmplete protease inhibitor cocktail (Roche), 1 mM EDTA and 1 mM or 3 mM TCEP (as indicated, when reducing condition was necessary for the experiment) were added and protein sample was kept in ice in the cold chamber until its use. When buffer exchange was needed it was performed with PD-10 desalting columns (Cytiva) previously equilibrated in the final desired buffer.

### Ellman’s assay

Ellman’s reagent, 5,5⍰-Dithiobis-(2-nitrobenzoic acid) (DTNB, Thermo Fisher Scientific), was dissolved in 20 mM sodium phosphate buffer pH 7.8, 150 mM NaCl. The buffer was previously purged with N_2_(g) before resuspending DTNB to a final concentration of 9 mM. Protein samples were desalted right before the assay using PD-10 desalting columns (Cytiva) equilibrated in Tris-HCl 20 mM, NaCl 150 mM, pH 8, to eliminate the reducing agent (when necessary). In each assay, the protein concentration was 10 µM and DTNB concentration was 50 µM, ensuring an excess of DTNB relative to the total number of cysteines in the sample (4 cysteine residues per VbrK^SD^ molecule); buffer was Tris-HCl 20 mM, NaCl 150 mM, pH 8. The mixture was incubated for 5 minutes at 25°C before measurement. UV-Visible absorption spectra were acquired on a Jasco V-530 spectrophotometer (Jasco, Easton, MD, USA). 2-nitro-5-thiobenzoate anion (TNB) concentration was determined from absorbance at 412 nm using a molar extinction coefficient of 14,150 M^-1^ cm^-1 35^.

### Circular Dichroism

Samples were desalted to 10 mM phosphate buffer, pH 7.8, 150 mM NaCl, with a PD-10 desalting column (Cytiva) and diluted to 10 µM final protein concentration in the same buffer. Spectra were acquired on a JASCO J-1500 spectropolarimeter from 190 to 260 nm at 50 nm/min scanning speed, with a data pitch of 0.1 nm, a bandwidth of 3 nm, an integration time of 1 s per point, and 8 scans were accumulated. Sample was in a quartz cuvette of 0.1 cm optical path length at 25 ºC. Data points for which the photomultiplier voltage was over 600 V were discarded.

### Cysteine blocking with 2-iodoacetamide reaction

100 µM VbrK^SD^ sample was mixed with locally synthesized 2-iodoacetamide at 50 mM final concentration, in 20 mM Tris-HCl, pH 8, 150 mM NaCl, and incubated for 2 hours at 25 ºC. The sample was centrifuged 5 minutes at 10,000 *g* and desalted with a PD-10 desalting column equilibrated in 5 mM Tris-HCl, pH 8, 150 mM NaCl.

### Ampicillin hydrolysis with NDM-1

A 6 mM ampicillin solution (300 µl) was hydrolyzed by addition of 0.5 µl of purified 500 µM NDM-1 β-lactamase and incubation for 10 min at room temperature. NDM-1 was inactivated adding 1 mM EDTA (final concentration).

### Nuclear magnetic resonance spectroscopy

Saturation transfer difference (STD) NMR spectroscopy experiments were acquired on a Bruker Avance III 700 MHz spectrometer equipped with a TXI probehead (Bruker, Billerica, MA, USA), employing standard Bruker stddiffesgp.3 pulse program. The protein saturation pulse was Sinc1.1000 with 50 ms duration. Saturation time was 2 s, and relaxation delay was 3 s. 4 repeats of 160 scans were accumulated. Off-resonance saturation frequency was -40 ppm and on-resonance saturation frequency was 0.75 ppm or 7 ppm. Temperature was 298 K. Protein and ligand concentrations were 20 µM and 400 µM respectively, in a 500 µl final volume. Buffer composition was 5 mM Tris-HCl, pH 8, 150 mM NaCl, 10% deuterium oxide. NMR data was processed and analyzed with TopSpin 4.5.

Heteronuclear single quantum coherence (^1^H-^15^N HSQC) experiments were conducted on a Bruker AVANCE II 800 MHz spectrometer equipped with a 5 mm TCI cryoprobe and an Avance Neo console, utilizing the standard Bruker pulse program *hsqcetfpf3gpsi*. Sweep widths were set to 12,500 Hz for ^1^H and 2,351 Hz for ^15^N, with 2,048 data points in the direct dimension (^1^H) and 128 increments in the indirect dimension (^15^N). The number of scans was 128, and the recycling delay was 1 second. NMR experiments were performed at 35 °C (308K) using 200 µM ^15^N-labeled VbrK^SD^ wild-type or 100 µM VbrK^SD^ C86A and C107A mutants, prepared by ^15^N enrichment with ^15^NH_4_Cl (Cambridge Isotope Laboratories) as the source of nitrogen in M9 minimal medium. Samples were prepared in 50 mM phosphate buffer, pH 7.8, 300 mM NaCl, and 7% (v/v) D_2_O. Reduced samples were preincubated with 3 mM TCEP for 10 minutes on ice.

### Protein Crystallization

VbrK^SD^ crystallization experiments were performed using the sitting-drop vapor diffusion method at 18 °C. Crystallization screening was conducted using a BCS screening kit in 96-well plates (Molecular Dimensions). For plate preparation, VbrK^SD^ was concentrated up to 7 mg/ml in a buffer containing 20 mM Tris-HCl, pH 8.0 and 150 mM NaCl. Plates were prepared with a Mosquito XTAL3 (SPT LABTECH), using 30 µl of reservoir solution, with 0.2 µl of protein mixed 1:1 with the reservoir solution.

Once optimal conditions were identified (0,2 M (NH_4_)_2_SO_4_, 0,1 M Tris-HCl, pH 8, 20% (w/v) PEG Smear Broad; Molecular Dimensions), crystals were grown by hanging-drop method in 48-well plates using 200 µl of crystallization solution. Drops consisted of 1.5 µL of protein solution mixed 1:1 with reservoir solution. For soaking, drops were treated with 1.5 µl of 9 mM TCEP solution to reach a final concentration of 3 mM for 10 min at 25 °C; at the same time, penicillin G powder was added.

Crystals were harvested at various incubation times up to 10 minutes, then soaked in a cryoprotectant solution containing 0.1 M (NH_4_)_2_SO_4_, 0.05 M Tris-HCl, pH 8, 10% (w/v) PEG Smear Broad, and 25% (v/v) glycerol. Crystals were flash-frozen in liquid N_2_ and shipped to the European Synchrotron Radiation Facility (ESRF).

### X-ray diffraction data collection and structure refinement

Diffraction data were collected at the European Synchrotron Radiation Facility (ESRF) at 100 K. Data were acquired by rotating the crystal in 0.1° increments up to a total rotation of 100° to minimize radiation damage. The diffraction data processing and scaling was performed with XDS ^36^.

The VbrK^SD^ structure was solved using molecular replacement using Phaser^38^ within the PHENIX suite^37^, using Protein Data Bank (https://www.rcsb.org/) ID 7CUS as the initial search model. The structure was completed in iterative rounds of manual model-building in COOT ^37^ and restrained refinement in phenix.refine ^38^. The quality of the final model was evaluated with phenix.molprobity ^39^. VbrK^SD^ crystallizes in space group R32, with one molecule per asymmetric unit. Density for all the polypeptide chain could be traced except for residue 23 in the N terminal and residues from 239 to 248 which include the C terminal TEV recognition sequence. There are no Ramachandran outliers and two strong densities in the solvent were modelled as SO_4_ ^-2^, present in the crystallization solution.

Diffraction data and model refinement statistics are shown in Table S2.

### Stopped-Flow Experiments

Changes in the visible spectrum of the chromogenic cephalosporin nitrocefin during the hydrolysis catalyzed by VbrK^SD^ were monitored using an Applied Photophysics SX18-MVR stopped-flow spectrometer equipped with a photodiode-array detector (Applied Photophysics, UK). Measurements were conducted in a buffer containing 20 mM Tris-HCl, pH 8, 150 mM NaCl, 1 mM TCEP, 1 mM EDTA, 1X cOmplete EDTA-free Protease Inhibitor Cocktail (Roche), at 6 °C. The data were adjusted to account for the instrument dead time of 2 ms. Spectral scans were recorded over a wavelength range of 190–730 nm, with an integration time of 10 ms and a 10 mm pathlength, 1000 time points were acquired over 1000 s. Nitrocefin solution in the syringe was 82 µM. VbrK^SD^ sample concentrations in the syringe were 82, 41 and 4 µM.

### Kinetic Analysis of Stopped-Flow Data

The reaction mechanism proposed for the reaction of VbrK^SD^ with β-lactams is shown in **Figure 8**.G. *k*_*1*_ was fixed to the diffusion-limited value of 10^8^ M^-1^ s^-1^ and the differential extinction coefficient for hydrolyzed free nitrocefin at 485 nm was 17,420 M^-1^ cm^-1 28^. Values of *k*_*−1*_, *k*_*2*_, *k*_*3*_ and the differential extinction coefficient for the enzyme-bound nitrocefin were obtained by non-linear least-squares minimization (minimize function from lmfit library in Python) between the experimental values of ΔAbsorbance at 485 nm vs. time and the integral of the differential equations system describing the species variations with time. The differential equations system was:

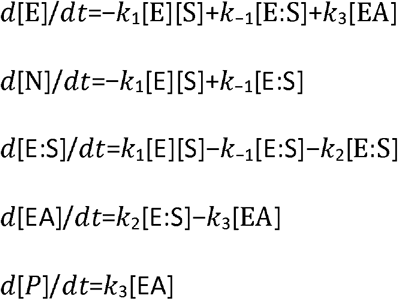

For the reaction of nitrocefin and the three different concentrations of VbrK^SD^ the parameters were optimized for the three curves simultaneously.

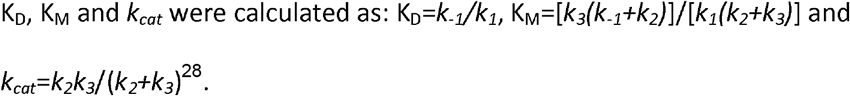

### Bocillin™-FL binding assays

Bocillin-FL (Invitrogen) was resuspended in absolute ethanol at a final concentration of 1 mM. 100 µM of VbrK^SD^ samples were mixed with 100 µM Bocillin-FL and incubated 20 min at 25 ºC. The reaction was stopped by adding trifluoroacetic acid (TFA) to 0,5% (v/v) final concentration and 5X Protein Sample Buffer (1X final concentration). An aliquot containing 3 µg of protein was loaded on each lane of a 16 % SDS-PAGE gel. Gels were visualized on a Typhoon FLA 7000 bioimager (GE Healthcare) with a FAM filter (Y520). After fluorescence detection, gels were stained with Coomassie Brilliant Blue.

### Cysteine labelling with DCIA

7-Diethylamino-3-[4’-(iodoacetamido)phenyl]-4-methylcoumarin (DCIA, Abcam ab145321) was resuspended in DMSO to a final concentration of 10 mg/mL (or 20,4 mM). TCEP was removed from TCEP-treated protein samples by buffer exchange to 20 mM Tris-HCl, 150 mM NaCl, pH 8 with PD10 desalting columns (Cytiva). 100 µM of desalted VbrK^SD^ samples were mixed with 500 µM DCIA and incubated ON at 4 ºC. 5X Protein Sample Buffer was added and 3 µg of protein were loaded on each lane of 16 % SDS-PAGE gels. Gels were visualized on a ChemiDoc XRS+ System (Bio-Rad) using the UV-transiluminator mode. After fluorescence detection, gels were stained with Coomassie Brilliant Blue.

### Intact Protein Mass Spectrometry and Top-Down analysis

Intact protein mass spectrometry analysis was performed in denaturing conditions. 100 µM protein samples were diluted 30 times with MilliQ water and 2 uL were injected for each sample. The separation was performed on LC instrument (Vanquish Horizon, Thermo Fisher Scientific) using Aquity UPLC Protein BEH C4 pre-column (2.1 x 5 mm, 1.7 µm, 300 A, Waters) with flow rate of 400 uL/min and oven temperature set to 45°C. The following gradient of solvent B was used: from 15 to 50 % within 3 min followed by column washing and re-equilibration steps. Solvent A was composed of MilliQ water with 0.1% formic acid, while solvent B consisted of acetonitrile with 0.1% formic acid. Eluting proteoforms were analyzed in positive polarity using Exploris 240 Orbitrap FT-MS instrument operated in the intact protein mode with Low Pressure option activated. Mass spectra were acquired as full scans in the range 700-4000 *m/z* at a resolution set to 120 000 or 240 000, RF Lens at 150 %, normalized AGC target of 300 %, maximum injection time of 200 ms, SID of 15 eV, while 10 µscans were averaged. The mass spectra were deconvolved using Biopharma Finder 5.3 (Thermo Scientific) with Xtract algorithm providing monoisotopic masse values.

To confirm the formation of covalent Penicillin G adduct and to localize the binding sites, samples of interest were submitted to top-down mass spectrometry analysis using higher energy collision induced dissociation (HCD). LC conditions were the same as the ones used for intact mass analysis in denaturing condition described above. Eluting proteoforms were analyzed on an Orbitrap Exploris 240 FT-MS benchtop instrument (Thermo Fisher Scientific, Bremen, Germany) operated in the intact protein mode with Low Pressure option activated, in positive polarity and using targeted MS/MS approach with ion multiplexing. For this, six precursor ions corresponding to different charge states of the same proteoform carrying Penicillin G adduct were input into the targeted inclusion mass list. Isolation window was set to 1.2 m/z, AGC target set as standard, maximum IT set as Auto, with 240’000 resolution at 200 m/z and averaging 10 µscans. Top-down mass spectrometry analysis was repeated using 4 different values (between 20-55 %) of normalized collision energy (NCE). Obtained Top-down data were processed using Peak-by-Peak software with MS deconvolution workflow (Spectroswiss, Lausanne, Switzerland).

### Western Blot

For the detection of VbrK^SD^(2-240), VbrK^SD^(25-240), and full length VbrK, the samples were resolved by 16% SDS-PAGE. The proteins were then transferred to a nitrocellulose membrane, using a solution of 200 mM Tris, 150 mM glycine, and 20% V/V methanol as transfer medium. For this, the TransBlot® TurboTM Blotting System (BIORAD) equipment was used, using the Standard SD protocol: 1 A, 25 V, 30 minutes. Then, the membrane was blocked for 17 h at 4 °C with shaking with 15 mL of a BLOTTO solution (100 mM Tris-HCl, pH 7.5; 150 mM NaCl, 3% skim milk powder, 3% BSA). Subsequently, four washes of 15 minutes each were performed with 15 mL of T-TBS (100 mM Tris-HCl, pH 7.5, 150 mM NaCl, 0.1% TWEEN 20), incubated with the rabbit IgG His-tag specific antibody conjugated to HRP (ab1187, Abcam) at a 1/50,000 dilution in T-TBS medium supplemented with 1% skim milk powder, gently stirring at room temperature for 1 h and then four more washes of 15 minutes each were performed with 15 mL of T-TBS. Finally, the membrane was developed by incubation with 1mL of the SuperSignal™ West Dura chemiluminescent reagent (Thermo Fisher Scientific) or with 1mL of the SuperSignal™ Pico chemiluminescent reagent (Thermo Fisher Scientific), according to the indications of the supplier. Chemiluminescent signals were captured either by exposure to CLXposure™ film (Thermo Fisher Scientific) followed by development with Kodak developer and fixation in Kodak fixer solution, or using a Bio-Rad ChemiDoc™ XRS+ Imaging System, as appropriate.

For penicillin-acylated VbrK^SD^ detection, Western blots were performed following the protocol described in Antinori *et al*.^40^ The membrane was then incubated with mouse anti-penicillin IgG monoclonal antibody (Pen9, Santa Cruz Biotechnology sc-57966) as primary antibody for 1 h at RT at a 1/100 dilution in T-TBS 1X + 2% BSA. Membranes were then incubated for 1 h at RT with a secondary antibody HRP-conjugated goat anti-mouse IgG (Abcam ab205719), at 1/10,000 dilution in T-TBS + 2% BSA washed 4 X 15 min with T-TBS 1X and developed using 1 mL SuperSignal™ West Dura chemiluminescent substrate (Thermo Fisher Scientific). Chemiluminescent signals were captured using a Bio-Rad ChemiDoc™ XRS+ Imaging System.

## Supporting information

Tables S1, S2 and S3. Figures S1-S14. The authors have cited additional references 26,41 within the Supporting Information.

## Supplementary Material Description

Tables S1, S2 and S3. Figures S1–S14. The authors have cited additional references^26,41^ within the Supporting Information.

## Author contributions

**Ignacio G. Palanca:** Investigation (lead); Formal Analysis (lead); Writing – Original Draft Preparation (supporting); Visualization (equal). **Irina P. Suárez:** Conceptualization (lead); Formal Analysis (lead); Funding acquisition (equal); Methodology (lead); Investigation (lead); Supervision (lead); Validation (lead); Visualization (lead); Writing – Original Draft Preparation (lead). **Maria J. Marcaida:** Methodology (equal); Formal Analysis (lead); Investigation (equal); Supervision (equal); Validation (lead); Writing – original draft (supporting). **Luciano A. Abriata:** Funding acquisition (supporting); Methodology (equal); Formal Analysis (equal). **Natalia Gasilova:** Mass Spectrometry analysis (lead); Writing – Original Draft Preparation (supporting). **Laure Menin:** Mass Spectrometry analysis (lead); Writing – Original Draft Preparation (supporting). **Franco E. Lacava:** Investigation (supporting). **Julieta Cairoli:** Investigation (supporting). **Matteo Dal Peraro:** Funding acquisition (supporting); Resources (equal). **Leticia I. Llarrull:** Conceptualization (lead); Validation (lead); Funding acquisition (lead); Writing – original draft (lead); Writing – review and editing (lead); Project administration (lead); Supervision (lead); Resources (lead).

## Acknowledgements

I.G.P. was a Ph.D. Fellow of Agencia Nacional de Promoción de la Investigación, el Desarrollo Tecnológico y la Innovación (Agencia I+D+i). I.G.P. was recipient of a Swiss Government Excellence Scholarship (ESKAS). I.G.P. is a Ph.D. fellow of CONICET. I.P.S. and L.I.L. are Staff members of CONICET. We thank Andrea Coscia, Alejandro Gago and Dr. Rodolfo M. Rasia from the Argentinean Structural Biology and Metabolomics Platform (PLABEM, IBR-UNR-CONICET) for their support on NMR experiments and Silvana Sut from IBR-UNR-CONICET for her technical assistance to research activities. We thank Dr. Sebastian A. Testero (IQUIR, CONICET-UNR, Argentina) for the synthesis of iodoacetamide. We thank the EPFL Protein Production and Structure Core Facility for providing the equipment for protein crystallization and the beam line scientists at ESRF beamline ID30a3 for support on diffraction data collection.

