## Supplementary material for "Redox-regulated cysteine acylation governs β-lactam sensing by the *Vibrio* histidine kinase VbrK": Tables S1, S2 and S3. Figures S1-S14. The authors have cited additional references 26,41 within the Supporting Information.

### Supplementary Tables

**Table S1.** Oligonucleotides used in this study. Restriction sites are in italics and underscored. Mismatches for site directed mutagenesis are bold and underscored.

| Experiment | Primer name | Sequence | Restriction site |
|---|---|---|---|
| Cloning of VbrK$^{SD}$-2-240 | OLL107 (fw) | 5'- CT*CCATGG*GGATCAAGCAGTTCCTGCTG -3' | NcoI |
| Cloning of VbrK$^{SD}$-25-240 | OLL108 (fw) | 5'- AT*CCATGG*GCCTGCCGGAACGTATC -3' | NcoI |
| Cloning of VbrK$^{SD}$ | OLL109 (rv) | 5'- A*GAATTC*GTGGTCCTCCACATCCC -3' | EcoRI |
| Replacement of thrombin site for TEV site | OLL178 (fw) | 5'- *GGATCC*GAAAACCTGTATTTTCAGGGCAGC*CTCGAG*CACC -3' | BamHI, XhoI |
| Replacement of thrombin site for TEV site | OLL179 (rev) | 5'- CTGAAAATACAGGTTTTC*GGATCC*GTGGTCCTCCACATCCCAG -3' | BamHI |
| C86A mutant | OLL257 (fw) | 5'- CCTGGCGAACACC**GCC**CGTGGTAAACTGC -3' | |
| C86A mutant | OLL258 (rev) | 5'- GCAGTTTACCACG**GGC**GGTGTTCGCCAGG -3' | |
| C107A mutant | OLL263 (fw) | 5'- CACCCGTGCGATT**GCC**AAGGGCACCC -3' | |
| C107A mutant | OLL264 (rev) | 5'- GGGTGCCCTT**GGC**AATCGCACGGGTG -3' | |
| L82A mutant | OLL236 (fw) | 5'- GTACAGC**GCA**GCGAACACCTGCCGTGGTA -3' | |
| L82A mutant | OLL237 (rev) | 5'- TTCGC**TGC**GCTGTACAGTTGCTGAATGTCCT -3' | |

**Table S2. Data collection and refinement statistics of the reduced VbrK$^{SD}$ structure; PDB code 28QS.** *Values in parentheses are for highest-resolution shell.

| | **Reduced VbrK$^{SD}$** |
|---|---|
| **Data collection** | |
| Space group | R32: H |
| **Cell dimensions** | |
| a, b, c (Å) | 98.0 98.0 164.6 |
| $\alpha$, $\beta$, $\gamma$ (º) | 90, 90, 120 |
| Resolution (Å) | 41.1-2.8 (2.97-2.8) |
| $R_{merge}$ (%) | 4.4 (49.4) |
| CC 1/2 | 1.0 (0.9) |
| $I/\sigma I$ | 24.4 (3.0) |
| Completeness (%) | 98.8 (99.5) |
| Redundancy | 5.7 (5.9) |
| **Refinement** | |
| Resolution (Å) | 41.1-2.8 |
| No. reflections | 7636 |
| $R_{work}/R_{free}$ | 0.24/0.27 |
| **No. atoms** | |
| Protein | 1737 |
| Ligand/ion | 10 |
| Water | 0 |

| | |
|---|---|
| Average B all atoms (Å$^2$) | 88.67 |
| **RMSD deviations** | |
| Bond length (Å) | 0.004 |
| Bond angles (º) | 0.77 |

**Table S3**. **Kinetic parameters for hydrolysis of nitrocefin catalyzed by VbrK$^{SD}$ wild-type (WT), C107A and L82A.** The kinetic parameters were obtained from the fit of the change of absorbance at 485 nm with time (data presented in Figure 8) to a system of differential equations corresponding to the reaction mechanism proposed in Figure 8.G. The dissociation constant $K_D$ would be around 490 µM for the WT protein, as calculated from the rate constants ($k_{+1}$ and $k_{-1}$); estimation is limited by the uncertainty in $k_{-1}$.

| Protein | $k_{-1}$ ($s^{-1}$) | $k_2$ ($s^{-1}$) | $k_3$ ($s^{-1}$) | $\Delta\varepsilon_{EA}$ ($M^{-1}$ $cm^{-1}$) |
|---|---|---|---|---|
| VbrK$^{SD}$ WT | (50 ± 130) $10^3$ | (1.0 ± 0.3) $10^{-3}$ | (2.50 ± 0.02) $10^{-3}$ | (70 ± 1) $10^3$ |
| C107A | (20 ± 100) $10^3$ | (0.6 ± 7.0) $10^{-3}$ | (1.9 ± 0.1) $10^{-3}$ | (50 ± 20) $10^3$ |
| L82A | (10 ±140) $10^3$ | (0.4 ± 5.0) $10^{-3}$ | (2.70 ± 0.02) $10^{-3}$ | (50 ± 20) $10^3$ |

#### Supplementary Figures

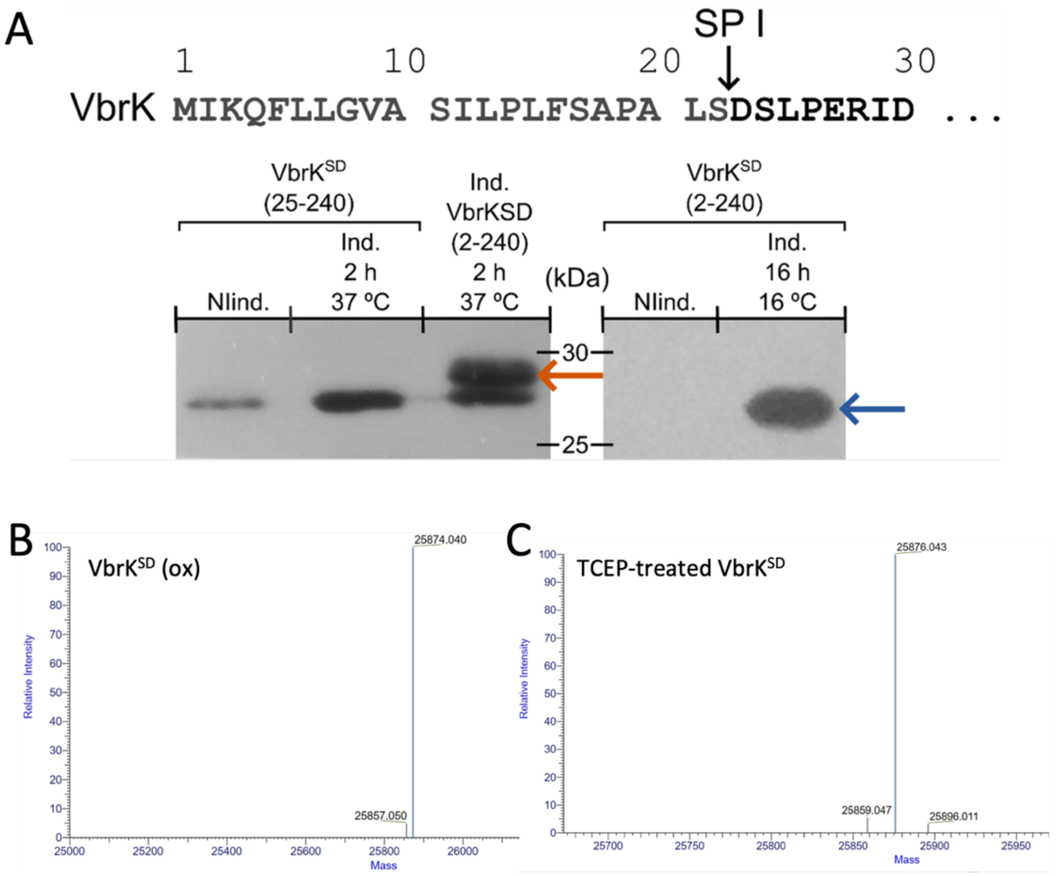


***Figure S 1. VbrK has a signal peptide that is cleaved between residues 22 and 23. A.*** *The presence of a signal peptide in VbrK was evaluated using the SignalP 6.0 server[1]. A potential signal peptide was detected in the N-terminal region of VbrK with a 0.97 probability of cleavage between residues 22 and 23. This signal peptide would facilitate the protein's translocation with the assistance of the Sec system.* ***A****. Top: sequence of the first 30 amino acids of VbrK, the arrow indicates the Signal Peptidase I (SP I) predicted cleavage site, between residues 22 and 23. Bottom: Western Blots of expression tests of* $VbrK^{SD}$*-(25-240)-6xH and* $VbrK^{SD}$*-(2-240)-6xH revealed with anti-Histidine-Tag-HRP conjugated antibody, orange arrow points at unprocessed* $VbrK^{SD}$*-(2-240) and blue arrow points at* $VbrK^{SD}$*-(2-240) after signal peptide cleavage. NInd.: non*

*induced culture, Ind.: IPTG-induced culture. Cultures at 37 ºC were induced with 500 µM IPTG, and culture at 16 ºC was induced with 10 µM IPTG. The best yields of soluble protein were obtained when the first 22 residues were included in the construct and sufficient time was given for full processing of the signal peptide: VbrK$^{SD}$ $_{2\text{-}240}$, induced with 10 µM IPTG for 16 h at 16 ºC. In this case, construct had a Thrombin site to remove the 6xH tag after affinity purification.* ***B****. Deconvolved mass spectrum of SEC-purified VbrK$^{SD}$ (after cleavage of 6xH-tag with TEV protease). Performed intact protein mass analysis detected the monoisotopic mass of 25874.0 Da (±0.1 Da), that matched, with 4.5 ppm mass accuracy, the theoretical mass of VbrK$^{SD}$ after cleavage of the signal peptide (residues 1-22) with the four cysteine residues engaged in two disulfide bonds.* ***C****. Deconvolved mass spectrum of TCEP-treated SEC-purified VbrK$^{SD}$ (after cleavage of 6xH-tag with TEV protease). Performed intact protein mass analysis detected the monoisotopic mass of 25876.0 Da (±0.1 Da), that matched, with 4.5 ppm mass accuracy, the theoretical mass of oxidized VbrK$^{SD}$ (B) with two additional protons, consistent with reduction of a disulfide bond.*

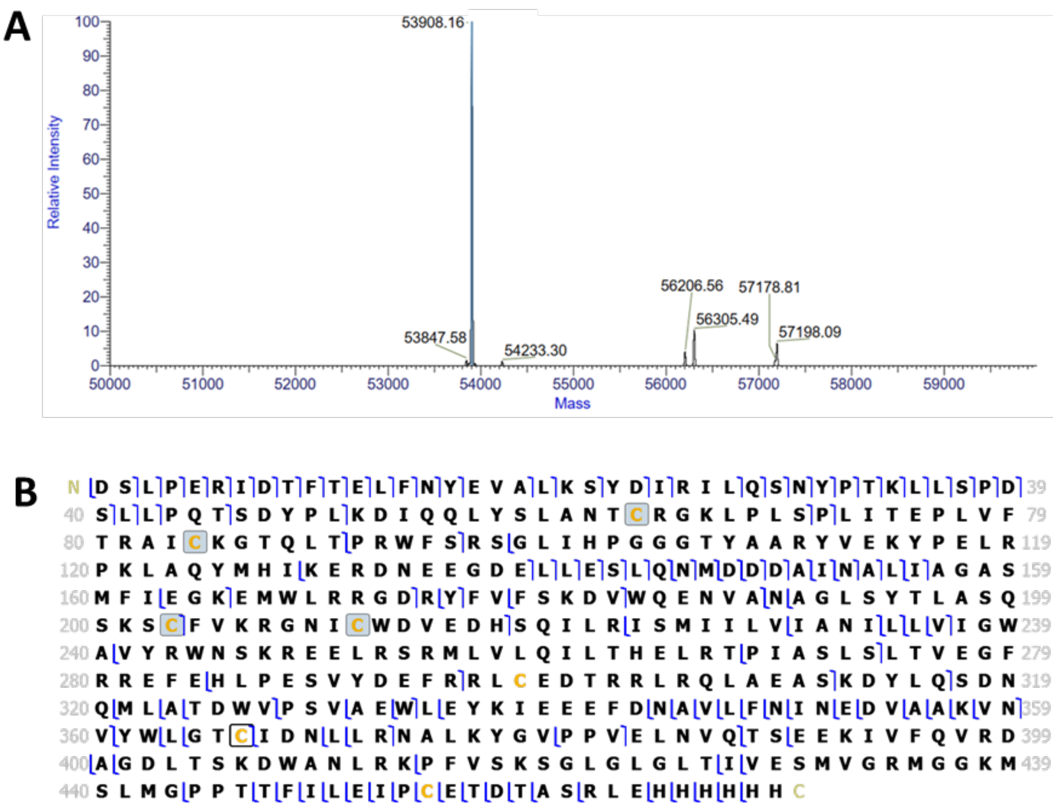


***Figure S 2. Intact protein and top-down mass spectrometry analyses of full-length VbrK*. A.** *Deconvolved mass spectrum of fully processed full length VbrK after cleavage of signal peptide at the same position as in the sensor domain construct, between residues 22 and 23. Performed intact protein mass analysis detected the average mass of 53908.2 Da (±0.4 Da), that matched the theoretical one with 7 ppm mass accuracy.* **B.** *Fragmentation map of full length VbrK obtained from top-down analysis by LC-MS/MS with 8 different normalized collision energies (NCEs) of HCD. Assigned characteristic b- and y-fragment ions are marked in blue. Grey boxes on Cys residues correspond to formation of disulfide bridges. Achieved sequence coverage of 32 % with 5 ppm mass accuracy for fragment assignment confirms the theoretical sequence for this protein.*

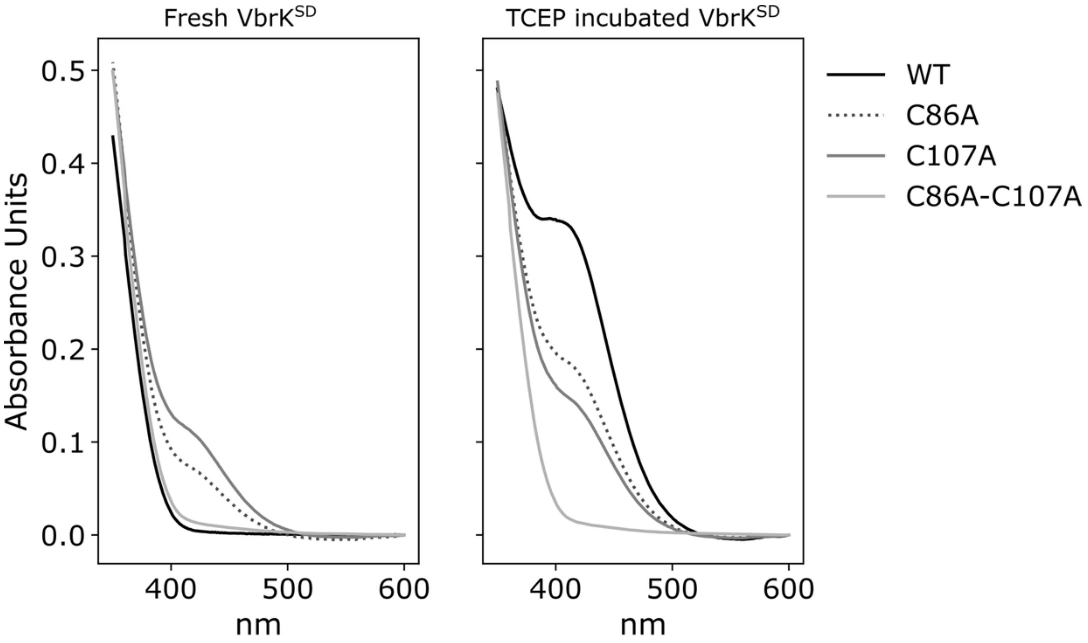


| | Abs 412 nm | [Reactive Cys] (µM) | Reactive Cys/molecule |
|---|---|---|---|
| VbrKSD WT (fresh) | 0.005 ± 0.004 | 0.4 ± 0.3 | 0.04 ± 0.03 |
| VbrKSD WT (TCEP Treated) | 0.296 ± 0.030 | 21 ± 2 | 2.1 ± 0.2 |
| VbrKSD C86A (fresh) | 0.096 ± 0.014 | 7 ± 1 | 0.7 ± 0.1 |
| VbrKSD C86A (TCEP Treated) | 0.156 ± 0.024 | 11 ± 2 | 1.1 ± 0.2 |
| VbrKSD C107A (fresh) | 0.111 ± 0.008 | 8 ± 1 | 0.8 ± 0.1 |
| VbrKSD C107A (TCEP Treated) | 0.134 ± 0.014 | 10 ± 1 | 1.0 ± 0.1 |
| VbrKSD C86A-C107A (fresh) | 0.017 ± 0.003 | 1.2 ± 0.2 | 0.12 ± 0.02 |
| VbrKSD C86A-C107A (TCEP Treated) | 0.048 ± 0.025 | 3 ± 2 | 0.3 ± 0.2 |

***Figure S 3. Ellman's assay on VbrK$^{SD}$ cysteine mutants suggests the disulfide bridge opening upon TCEP incubation is C86-C107.*** *UV-Visible absorbance spectra of 10 µM of VbrK$^{SD}$ WT, C86A, C107A, and C86A-C107A before (left) and after (right) incubation with TCEP, in the presence of 50 µM DTNB. The spectra shown correspond to one Ellman's assay. The corresponding calculations of Cys concentration and Reactive Cys/molecule for that experiment are shown in the table below. Error from Ellman's assay correspond to three independent protein purifications and thiol and protein quantifications.*

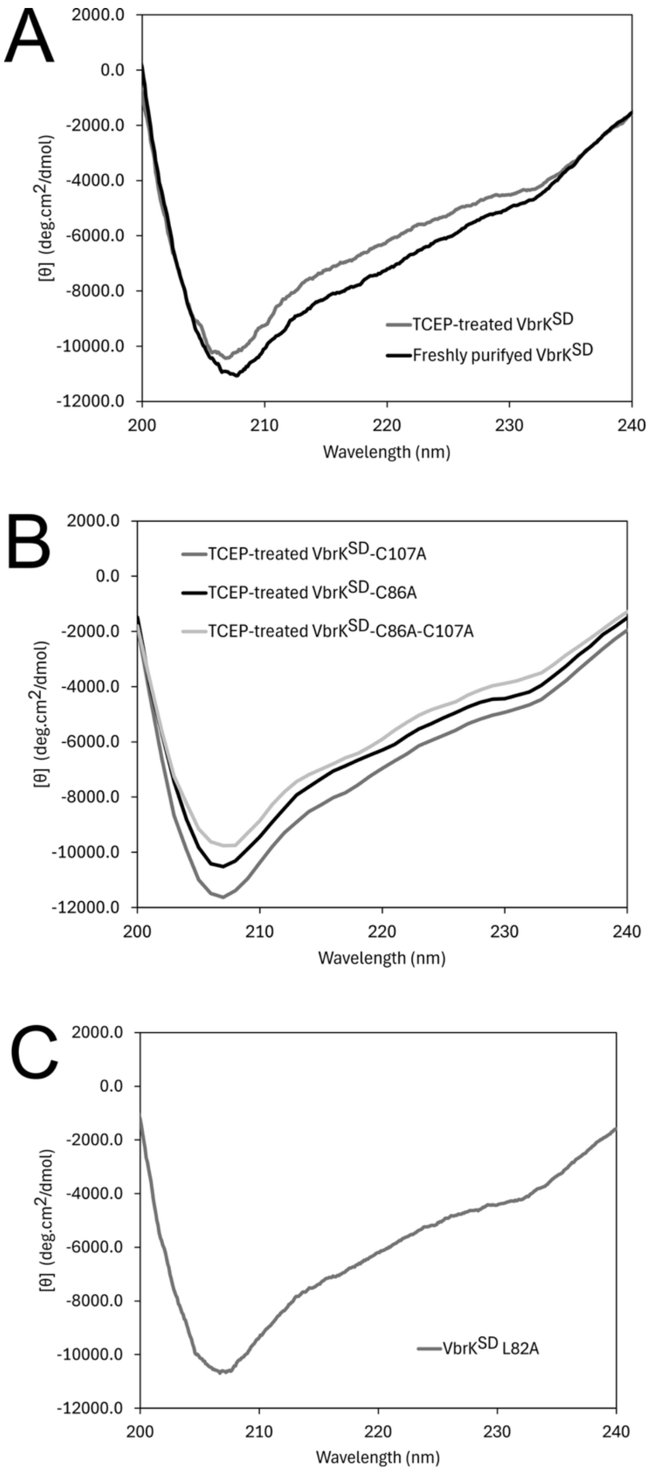


***Figure S 4. Circular dichroism spectra of oxidized and reduced VbrK$^{SD}$ wild-type and the C86A, C107A, C86A-C107A and L82A mutants. A.*** *Circular dichroism spectra of freshly purified VbrK$^{SD}$ WT and of VbrK$^{SD}$ WT after TCEP treatment.* ***B.*** *Circular dichroism spectra of TCEP-treated VbrK$^{SD}$ C86A, C107 A and C86A-C107A.* ***C.*** *Circular*

*dichroism spectra of freshly purified VbrK$^{SD}$ L82A. In all cases, protein concentration was 10 µM.*

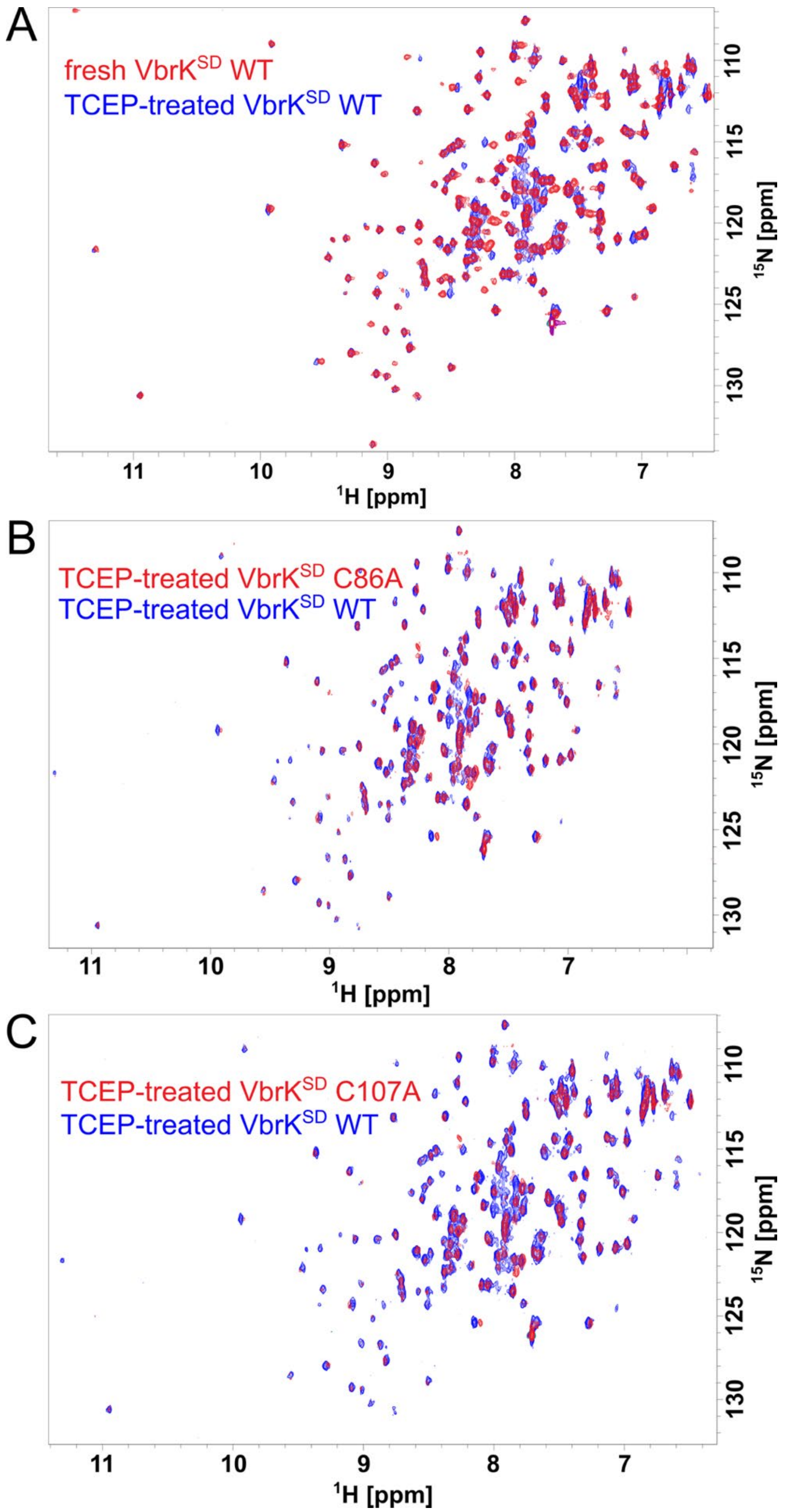


***Figure S 5. The $^{1}H$-$^{15}N$ HSQC spectra of wild-type VbrK$^{SD}$ and of the C86A and C107A mutants revealed reduction of the C86-C107 disulfide bond upon TCEP-treatment.*** *Full*

*Spectra corresponding to Figure 3.* ***A.*** *2D $^{1}H$-$^{15}N$ HSQC spectrum of 200 µM $^{15}N$-labeled VbrK$^{SD}$ TCEP-treated (blue) and freshly purified (oxidized, red).* ***B.*** *2D $^{1}H$-$^{15}N$ HSQC spectrum of 200 µM $^{15}N$-labeled TCEP-treated VbrK$^{SD}$ WT (blue) and 100 µM TCEP-treated C86A variant (red).* ***C.*** *2D $^{1}H$-$^{15}N$ HSQC spectrum of 200 µM $^{15}N$-labeled TCEP-treated VbrK$^{SD}$ WT (blue) and 100 µM TCEP-treated C107A variant (red).*

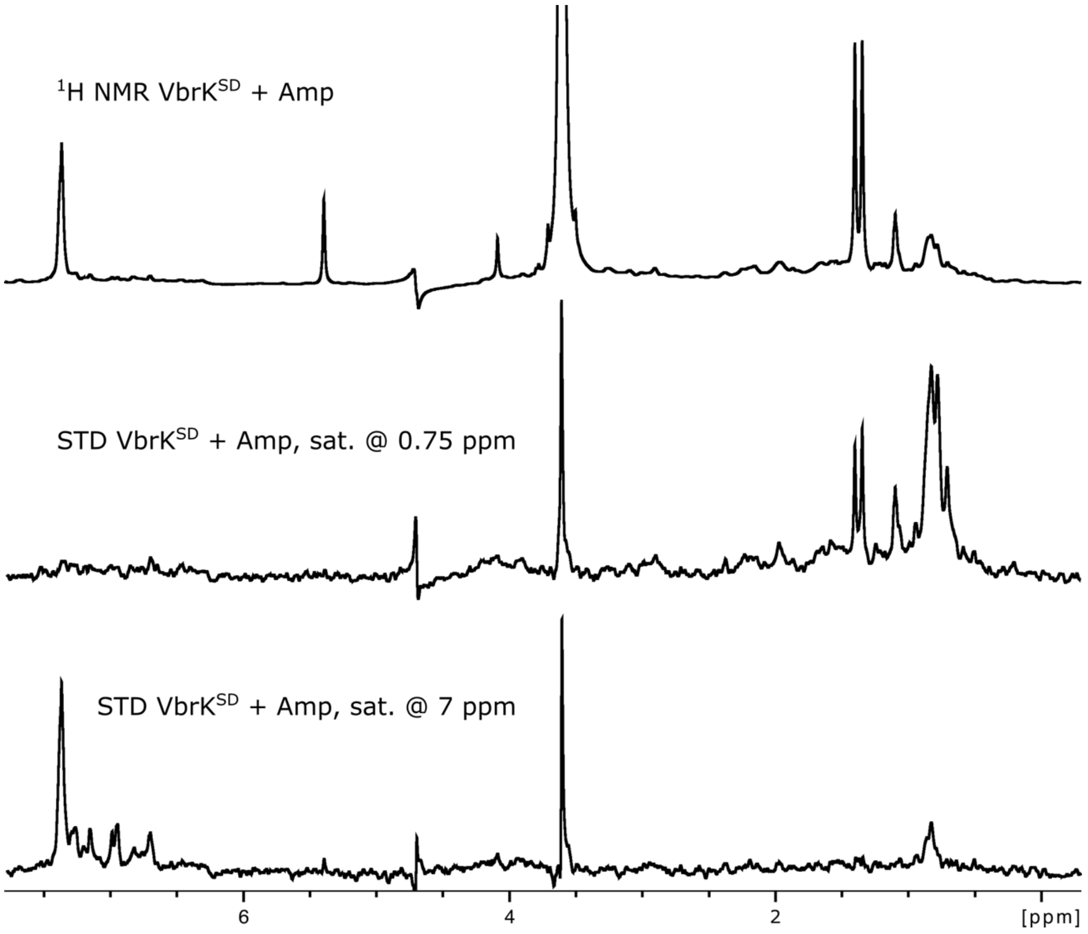


***Figure S 6. Oxidized VbrK$^{SD}$ does not interact with ampicillin, even after storage in buffer with no reducing agent, demonstrating the stability of the disulfide bonds in the purified sensor domain.*** *STD NMR experiments were carried out saturating at 0.75 and 7 ppm (on resonance). VbrK$^{SD}$ was incubated on ice for 66 h at 4 °C in the absence of TCEP, to compare with spectra shown in Figure 1. Buffer was 5 mM Tris-HCl, 150 mM NaCl, 10% deuterium oxide, pH 8. From top to bottom: $^1$H NMR spectrum of 20 µM VbrK$^{SD}$ with 400 µM ampicillin; STD spectrum of the same sample saturating at 0.75 ppm; STD spectrum of the same sample saturating at 7 ppm.*

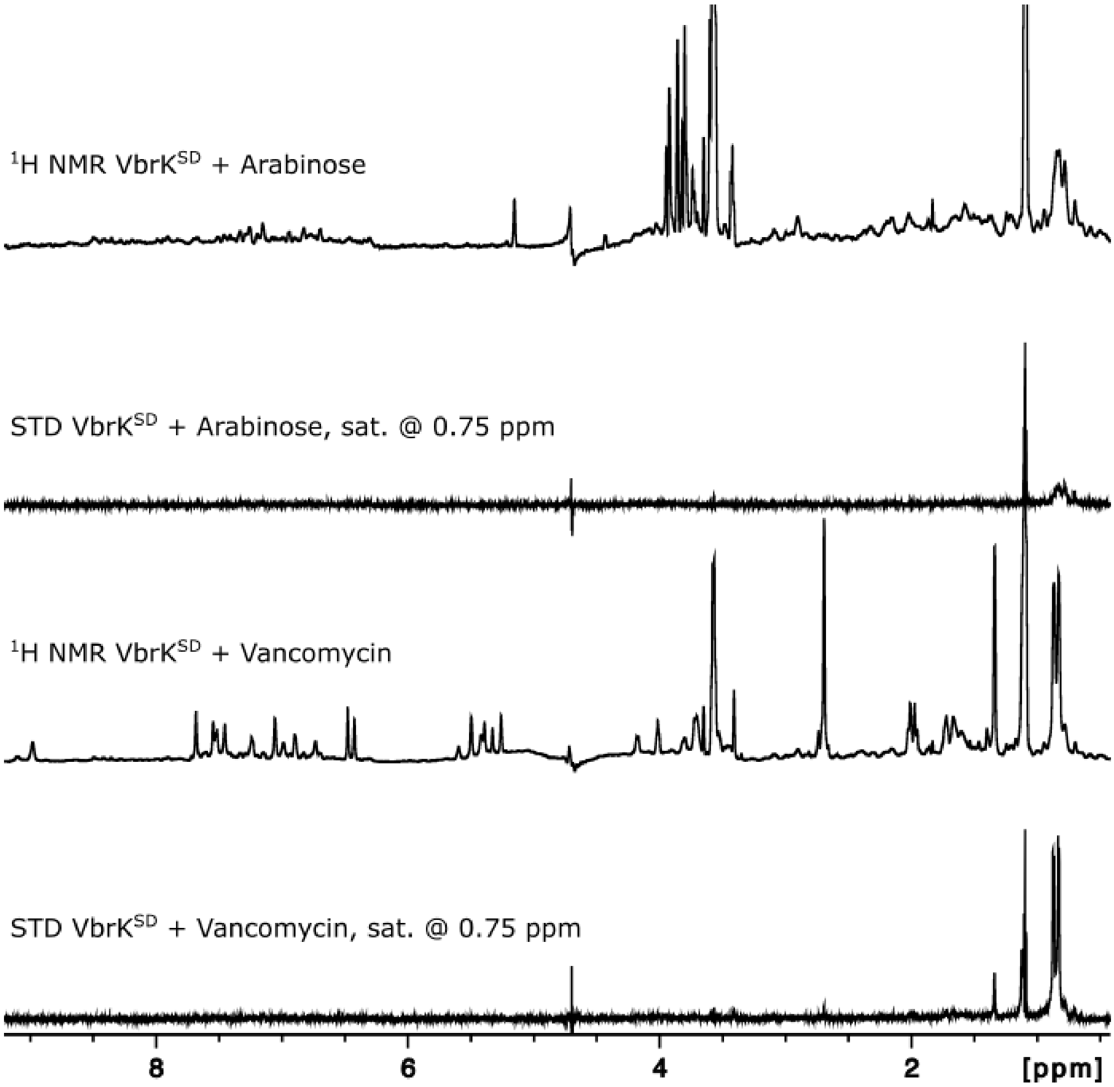


***Figure S 7. TCEP-treated VbrK$^{SD}$ does not interact with arabinose or vancomycin.*** *STD NMR experiments were carried out saturating at 0.75 ppm (on resonance). Buffer was 5 mM Tris-HCl, pH 8, 150 mM NaCl, 10% deuterium oxide. From top to bottom: $^{1}$H NMR spectrum of 20 µM TCEP-treated VbrK$^{SD}$ with 400 µM L-(+)-arabinose; STD NMR spectra of the same sample saturating at 0.75 ppm; $^{1}$H NMR spectrum of 20 µM TCEP-treated VbrK$^{SD}$ with 400 µM vancomycin; STD NMR experiment of the same sample saturating at 0.75 ppm.*

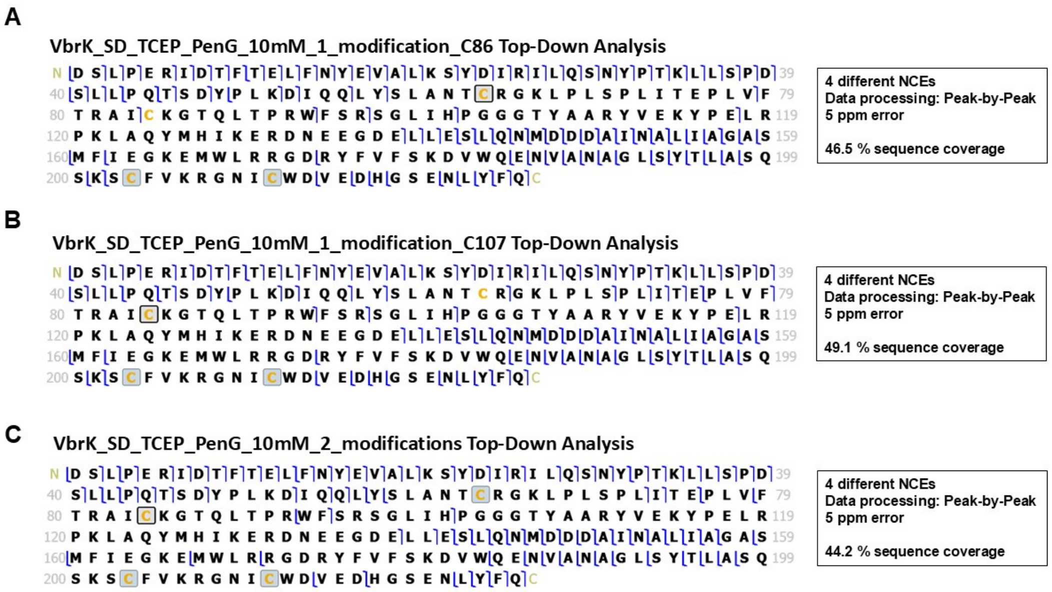


***Figure S 9. Top-down mass spectrometry analysis of VbrK$^{SD}$-PenG adducts identifies C86 and C107 as the sites of β-lactam acylation. A.*** *Fragmentation map of mono-PenG-acylated VbrK$^{SD}$ proteoform with PenG adduct on C86.* ***B.*** *Fragmentation map of mono-PenG-acylated VbrK$^{SD}$ proteoform with PenG adduct on C107.* ***C.*** *Fragmentation map of di-PenG-acylated VbrK$^{SD}$ proteoform with PenG adduct on both C86 and C107. Reaction conditions: 100 µM TCEP-treated VbrK$^{SD}$ incubated with 10 mM Penicillin G for 6 h at 25 °C. These top-down data were obtained by LC-MS/MS with 4 different HCD normalized collision energies (NCEs) applied for mono- and di-PenG-acylated proteoforms. One PenG adduct provided the mass shift of 334.0984 Da. Assigned characteristic b- and y-fragment ions are marked in blue. Grey boxes on C226 and C234 residues correspond to formation of a disulfide bridge. Corresponding sequence coverages achieved with 5 ppm mass accuracy are displayed for each proteoform. Intact mass spectra of all these samples are shown in Figure 7.*

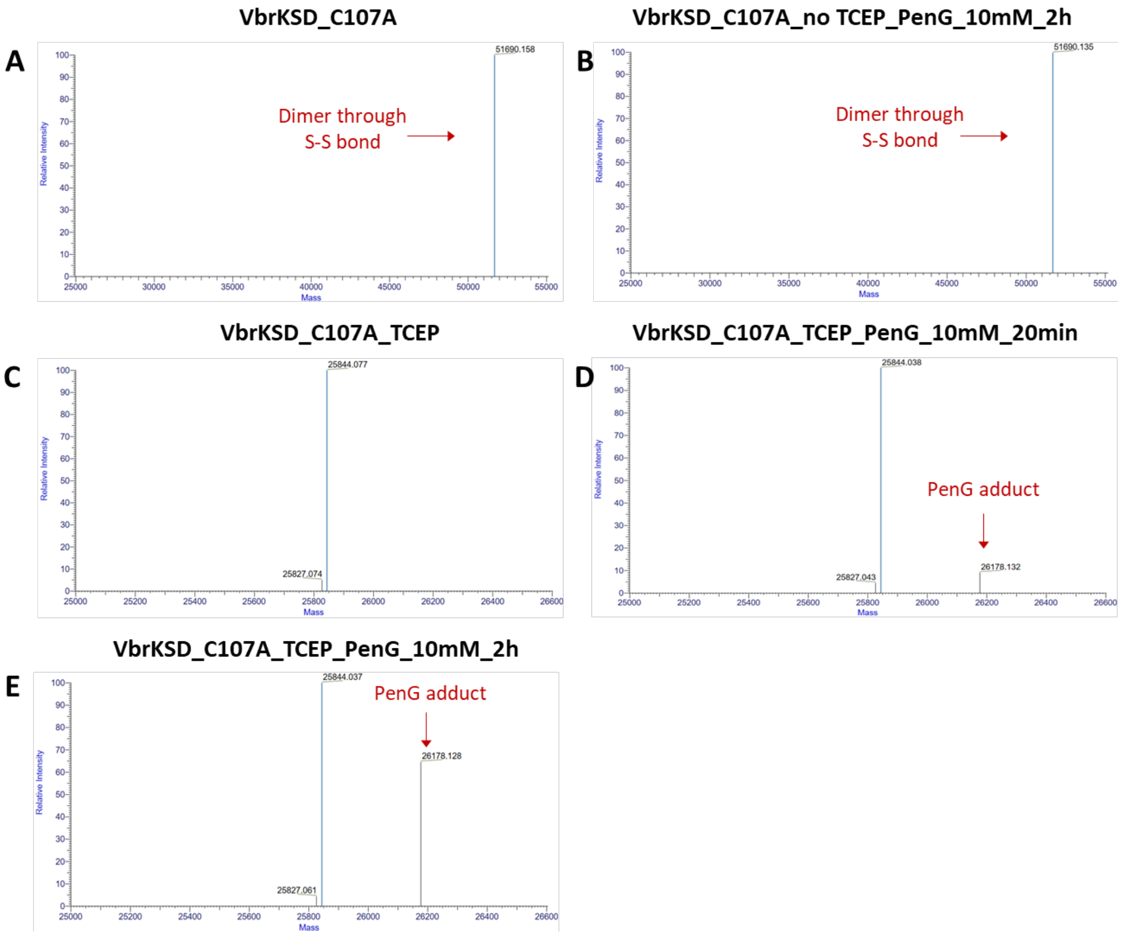


***Figure S 10. Intact protein mass spectrometry analysis with VbrK$^{SD}$ C107A shows Penicillin G-acylation on C86. A.*** *Deconvolved mass spectrum of VbrK$^{SD}$ C107A after storage at 4 °C and later -20 °C (100 µM); formation of a dimer through disulfide bond was detected.* ***B.*** *Deconvolved mass spectrum of VbrK$^{SD}$ C107A (100 µM; no TCEP treatment – same sample in A), incubated with 10 mM Penicillin G for 2 h at 25 °C.* ***C.*** *Deconvolved mass spectrum of TCEP-treated VbrK$^{SD}$ C107A (100 µM).* ***D.*** *Deconvolved mass spectrum of TCEP-treated VbrK$^{SD}$ C107A (100 µM) incubated with 10 mM Penicillin G for 20 min at 25 °C.* ***E.*** *Deconvolved mass spectrum of TCEP-treated VbrK$^{SD}$ C107A (100 µM) incubated with 10 mM Penicillin G for 120 min at 25 °C. A-E display the monoisotopic mass values in Da with mass accuracy of 5 ppm.*

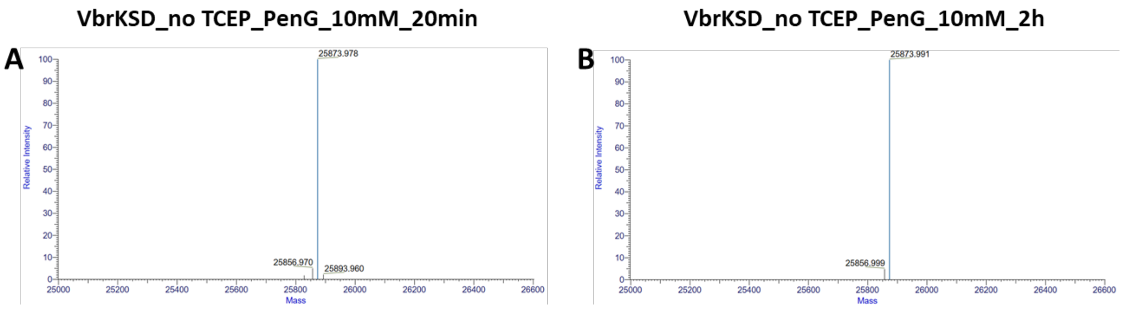


***Figure S 11. Intact protein mass spectrometry analysis with oxidized VbrK$^{SD}$ (no TCEP treatment) reveals no reaction with Penicillin G. A.*** *Deconvolved mass spectrum of oxidized VbrK$^{SD}$ (100 µM) incubated with 10 mM Penicillin G for 20 min at 25 °C.* ***B.*** *Deconvolved mass spectrum of oxidized VbrK$^{SD}$ (100 µM) incubated with 10 mM Penicillin G for 120 min at 25 °C. A and B display the monoisotopic mass values in Da with mass accuracy of 5 ppm.*

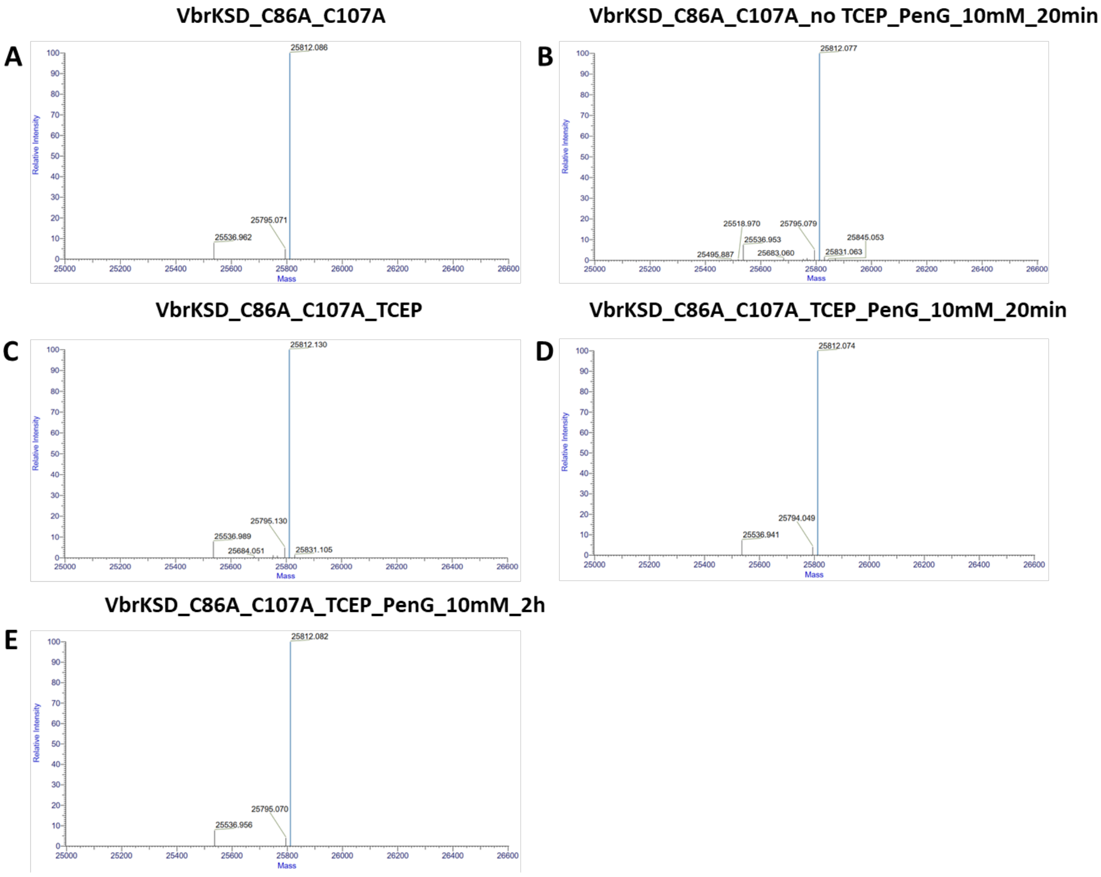


***Figure S 12. Intact protein mass spectrometry analysis with VbrK$^{SD}$ C86A C107A reveals no reaction with Penicillin G. A.*** *Deconvolved mass spectrum of VbrK$^{SD}$ C86A C107A (no TCEP treatment) after storage at 4 °C and later -20 °C (100 µM).* ***B.*** *Deconvolved mass spectrum of VbrK$^{SD}$ C86A C107A (85 µM; no TCEP treatment – same sample in A), incubated with 10 mM Penicillin G for 20 min at 25 °C.* ***C.*** *Deconvolved mass spectrum of TCEP-treated VbrK$^{SD}$ C86A C107A (85 µM).* ***D.*** *Deconvolved mass spectrum of TCEP-treated VbrK$^{SD}$ C86A C107A (85 µM) incubated with 10 mM Penicillin G for 20 min at 25 °C.* ***E.*** *Deconvolved mass spectrum of TCEP-treated VbrK$^{SD}$ C86A C107A (85 µM) incubated with 10 mM Penicillin G for 120 min at 25 °C. A-E display the monoisotopic mass values in Da with mass accuracy of 5 ppm.*

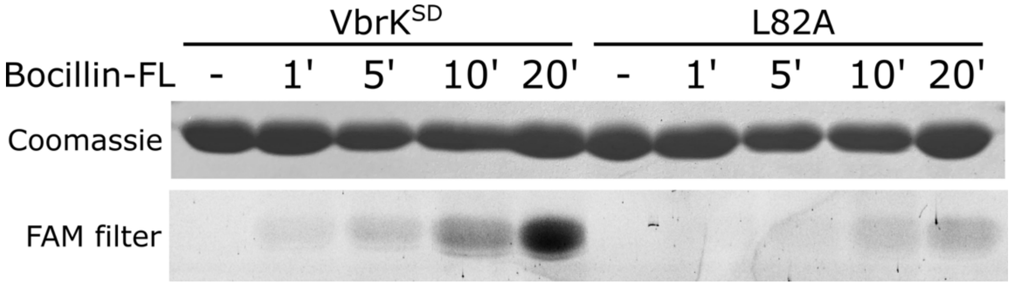


***Figure S 13. VbrK$^{SD}$ L82A fails to accumulate the covalent adduct with Bocillin-FL.*** *100 µM TCEP-treated VbrK$^{SD}$ WT and L82A were incubated with 100 µM Bocillin-FL for 20 minutes at 25 ºC. Aliquots were taken at different time points and the reaction was stopped by adding 0.5% TFA and 1X protein sample buffer. Samples were run on a 16% SDS-PAGE and the fluorescent bands were detected on a Typhoon™ FLA 7000 with a FAM filter prior to Coomassie Brilliant Blue staining.*

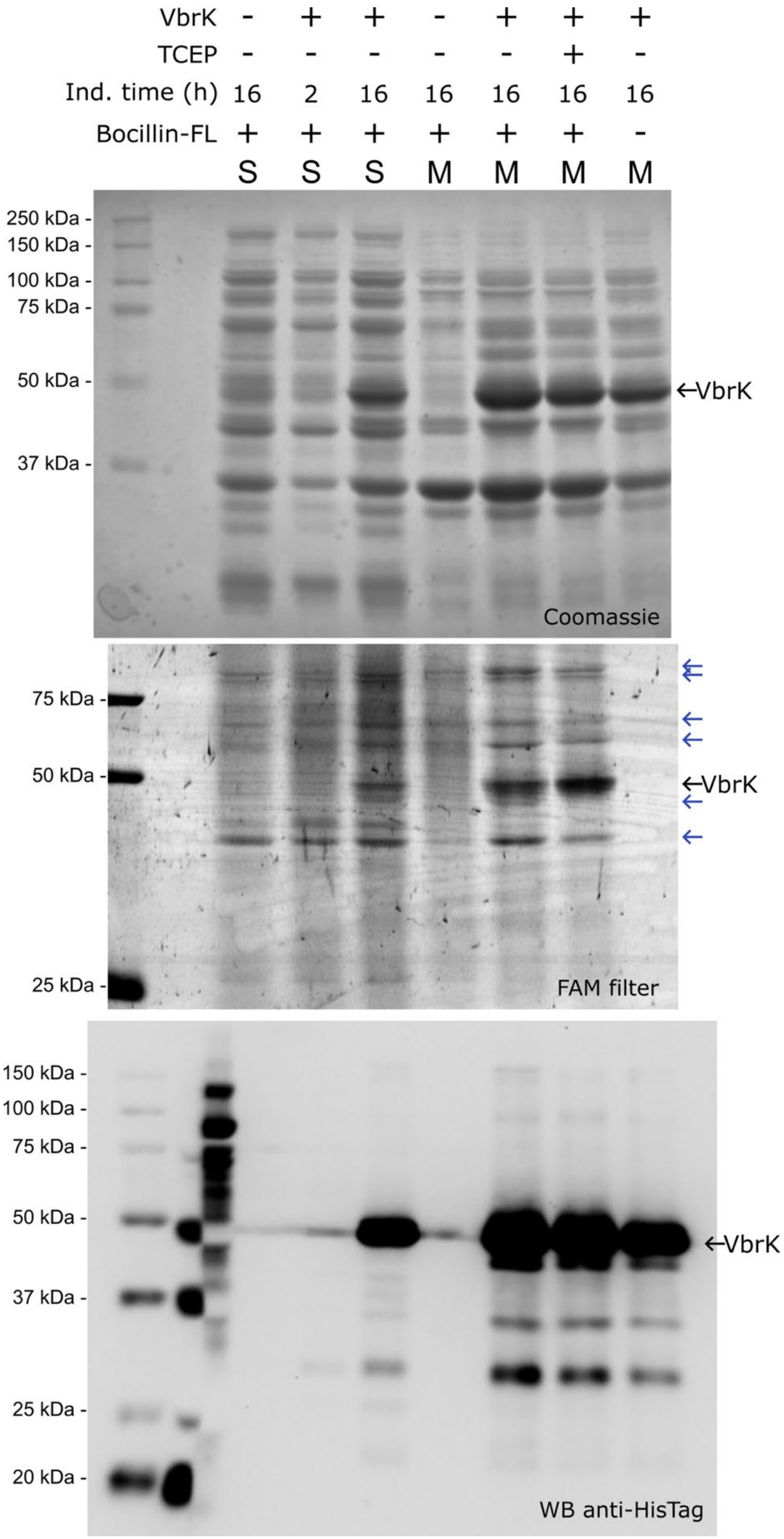


***Figure S 14. Intrinsic reduction of VbrK in cellular membrane contexts enables covalent Bocillin-FL labeling.*** *Complete gels and Western blot shown in Figure 9. Full-*

*length VbrK was labeled with Bocillin-FL in E. coli BL21 Star™ DE3 whole cells, spheroplasts (S), and membrane protein extracts (M), followed by analysis on a 16% SDS-PAGE gel. Expression of VbrK in E. coli BL21 Star™ DE3 was induced with 10 µM IPTG during 2 or 16 h. Labeling with Bocillin-FL in whole cells and spheroplasts was performed by incubation with 80 µM Bocillin-FL (final concentration) for 20 min at 25°C. Membrane protein preparation from E. coli cells expressing VbrK were incubated with Bocillin-FL without or with pre-treatment with TCEP. Top image: gel where all proteins were stained with Coomassie Brilliant Blue. Middle image: same gel as on top before Coomassie staining, where Bocillin-FL-labelled (fluorescent) proteins were detected by scanning the gel in a Typhoon FLA-7000 equipment, using the FAM filter. Bottom image: Western blot using an α-His-Tag-HRP antibody (ab1187, Abcam; dilution 1/25000) to detect the C-terminal His-tag (6xH) in recombinant VbrK. The section of the membrane with the Precision Plus Protein WesternC™ Blotting Standards (1/5 dilution) was cut and incubated with Streptactin-HRP (Lane 1). Dual Color Standards marker was used to guide cutting of the membrane (Lane 2). VbrK (ca. 54 kDa; black arrow) and the five E. coli PBPs were labeled with Bocillin-FL in spheroplasts and membrane extracts. The E. coli PBPs (blue arrows) detected were PBP1a/1b (~94 kDa); PBP2 (65 kDa); PBP3 (58 kDa); PBP4 (50 kDa); PBP5 (43 kDa), as previously reported[2]. VbrK was labelled both in spheroplasts and membranes, without requirement of treatment with reducing agents. Lane 1: Precision Plus Protein WesternC™ Blotting Standards (Bio-Rad). Lane 2: empty or Precision Plus Protein Dual Color Standards (Bio-Rad) in WB gel. Lane 3: spheroplasts prepared from E. coli cells transformed with empty pET24a(+), induced with IPTG 10 µM for 16 h. Lanes 4 and 5: spheroplasts prepared from E. coli cells transformed with pET24a(+)_vbrK induced with*

*10 µM IPTG for 2 h (4), and 16 h (5), and then incubated with 80 µM Bocillin-FL for 20 min at 25°C. Lane 6: Membrane protein preparation from E. coli cells transformed with empty pET24a(+), induced with IPTG 10 µM for 16 h. Lanes 7 to 9: Membrane protein preparation from E. coli cells transformed with pET24a(+)_vbrK induced with 10 µM IPTG for 16 h, incubated with 80 µM Bocillin-FL for 20 min at 25°C after preparation (7), after treatment with 3 mM TCEP for 10 minutes (8), and without incubation with Bocillin-FL (9).*
